# Host range of naturally and artificially evolved symbiotic bacteria for a specific host insect

**DOI:** 10.1101/2024.04.28.591543

**Authors:** Ryuga Sugiyama, Minoru Moriyama, Ryuichi Koga, Takema Fukatsu

## Abstract

Diverse insects are intimately associated with specific symbiotic bacteria, where host and symbiont are integrated into an almost inseparable biological entity. These symbiotic bacteria usually exhibit host specificity, uncultivability, reduced genome size, and other peculiar traits relevant to their symbiotic lifestyle. How host-symbiont specificity is established at the very beginning of symbiosis is of interest but poorly understood. To gain insight into the evolutionary issue, we adopted an experimental approach using the recently developed evolutionary model of symbiosis between the stinkbug *Plautia stali* and *Escherichia coli*. Based on the laboratory evolution of *P. stali-E. coli* mutualism, we selected Δ*cyaA* mutant of *E. coli* as an artificial symbiont of *P. stali* that has established mutualism by a single mutation. In addition, we selected a natural cultivable symbiont of *P. stali* of relatively recent evolutionary origin. These artificial and natural symbiotic bacteria of *P. stali* were experimentally inoculated to symbiont-deprived newborn nymphs of diverse stinkbug species. Strikingly, the mutualistic *E. coli* was unable to establish infection and support growth and survival of all the stinkbug species except for *P. stali*, uncovering that host specificity can be established at a very early stage of symbiotic evolution. Meanwhile, the natural symbiont was able to establish infection and support growth and survival of several stinkbug species in addition to *P. stali*, unveiling that a broader host range of the symbiont has evolved in nature. Based on these findings, we discuss what factors are relevant to the establishment of host specificity in the evolution of symbiosis.

**IMPORTANCE:** How does host-symbiont specificity emerge at the very beginning of symbiosis? This question is difficult to address, because it is generally difficult to directly observe the onset of symbiosis. However, recent development of experimental evolutionary approaches to symbiosis has brought about a breakthrough. Here we tackled this evolutionary issue using a symbiotic *Escherichia coli* created in laboratory and a natural *Pantoea* symbiont, which are both mutualistic to the stinkbug *Plautia stali*. We experimentally replaced essential symbiotic bacteria of diverse stinkbugs by the artificial and natural symbionts of *P. stali*, and evaluated whether the symbiotic bacteria evolved for a specific host can establish infection and support growth and survival of heterospecific hosts. Strikingly, the artificial symbiont showed strict host specificity to *P. stali* whereas the natural symbiont was capable of symbiosis with diverse stinkbugs, which provide insight into how host-symbiont specificity can be established at early evolutionary stages of symbiosis.

## INTRODUCTION

Many insects are obligatorily associated with their specific bacterial symbionts, where the host insects cannot survive without their microbial partners (1–3). At the beginning, such obligatory symbiotic bacteria must have been derived from environmental free-living bacteria. Understanding how originally unrelated host and symbiont have established symbiosis, developed interdependence, and become integrated into a coherent biological entity is an intriguing issue in the field of evolutionary biology (4, 5).

In highly mutualistic associations such as aphid-*Buchnera* and tsetse-*Wigglesworthia* symbioses (6, 7), the host phylogeny generally mirrors the symbiont phylogeny, reflecting stable vertical symbiont transmission and consequent host-symbiont co-speciation over evolutionary time (8, 9). During the course of co-evolution, the bacterial symbionts tend to develop dependence on their hosts, lose the capability of independent life, become uncultivable, suffer genome reduction, etc., thereby constituting a highly integrated symbiotic system where neither the host nor the symbiont can thrive independently (10, 11). Here, each host lineage is consistently associated with its own specific symbiont lineage, where high host-symbiont specificity is likely to evolve (12, 13). On the other hand, among facultative symbiotic associations such as *Wolbachia*, *Serratia*, *Sodalis,* etc. with diverse insects (14–16), or even among evolutionarily conserved mutualistic associations such as *Riptortus-Caballeronia* symbioses (17, 18), the bacterial symbionts are occasionally or constantly moving around different host lineages, where they tend to remain less dependent on their hosts, often retain the capability of surviving outside their hosts, remain cultivable in some cases, keep large genome sizes, etc., which will end up with such evolutionary consequences as host-symbiont phylogenetic promiscuity and compromised/context-dependent host-symbiont interdependence and specificity (13, 19). Theoretical as well as empirical studies have shed light on what factors may affect the evolutionary trajectories toward diverse symbiotic associations ranging from loose/exchangeable ones to strict/specific ones (20, 21).

How are host-symbiont interdependence and specificity established at the very beginning of the evolution of symbiosis? Needless to say, this question is difficult to address, because it is difficult to observe the onset of symbiotic evolution in most cases. However, recent development of experimental evolutionary approaches to symbiosis has opened a window to look into this unexplored research area (22, 23). Here we tackle this evolutionary issue by experimentally comparing an artificial insect symbiont that was created in laboratory with a natural insect symbiont that has evolved in nature.

The brown-winged green stinkbug *Plautia stali* (Hemiptera: Pentatomidae) has a specialized symbiotic organ consisting of numerous crypts in a posterior region of the midgut, in which a specific bacterial symbiont of the genus *Pantoea* densely populates (24). The symbiont is essential for growth and survival of the host insect and vertically transmitted to newborn nymphs by maternal smearing of symbiont-containing excrement onto eggshell (24, 25). Notably, the essential symbiotic bacteria in natural populations of *P. stali* exhibit geographic polymorphism across the Japanese Archipelago: an uncultivable symbiont *Pantoea* sp. A (= Sym A) is fixed in the main islands, whereas an uncultivable symbiont *Pantoea* sp. B (= Sym B) coexists with cultivable symbionts *Pantoea* sp. C-F (= Sym C-Sym F) in the southwestern Ryukyu islands (24). Hence, both uncultivable and cultivable bacterial strains are available as essential natural symbionts of *P. stali*, which are useful for a variety of experimental purposes (24). In this study, we used a Sym C strain as a natural cultivable symbiont of *P. stali*.

Recently, we established an experimental symbiotic system between *P. stali* as host and the model bacterium *Escherichia coli* as symbiont (26). We orally inoculated a hypermutating *E. coli* strain to symbiont-free newborn nymphs of *P. stali*, obtained a few adult insects that managed to survive and attained adulthood, dissected and homogenized the *E. coli*-populated symbiotic organ from each of them, inoculated the homogenate to symbiont-free newborn nymphs in the same way, and repeated the process for many host generations. Finally, we successfully obtained mutualistic evolutionary *E. coli* strains that can support growth, survival and reproduction of *P. stali*, and identified that single gene mutations disrupting the carbon catabolite repression global transcriptional regulator system for bacterial metabolic switching, namely Λ′*cyaA* and Λ′*crp*, make *E. coli* mutualistic to *P. stali* (26). Therefore, by introducing Λ′*cyaA* or Λ′*crp* mutation into the wild-type genetic background of *E. coli*, we can create a mutualistic *E. coli* strain artificially (26). In this study, we selected and used a Λ′*cyaA E. coli* mutant as a mutualistic *E. coli* strain.

Besides *P. stali*, diverse stinkbugs are also obligatorily associated with specific symbiotic bacteria of beneficial nature, which are similarly harbored in the midgut symbiotic organ and mostly, if not always, vertically transmitted to offspring via egg surface contamination or other means (27, 28). In this study, we established stable laboratory strains of diverse stinkbugs representing 7 species, 3 families and 2 superfamilies, generated newborn nymphs deprived of their original symbiont, inoculated the mutualistic *E. coli* strain Λ′*cyaA* or the natural symbiont strain Sym C to them, and investigated whether these artificial and natural symbiotic bacteria of *P. stali* can establish symbiosis with the different host species and what phenotypic consequences are observed with the replacement of their original symbiont by the artificial and natural symbionts of *P. stali*.

## RESULTS

### Establishment of stinkbug strains, removal of original symbiotic bacteria, and inoculation of natural and artificial symbiotic bacteria from *P. stali*

We have established stable laboratory strains of the following diverse stinkbug species: *P. stali*, *Glaucias subpunctatus*, *Nezara viridula* and *Halyomorpha halys* representing the family Pentatomidae; *Lampromicra miyakonus*, *Poecilocoris lewisi* and *Eucorysses grandis* representing the family Scutelleridae; and *Riptortus pedestris* representing the family Alydidae (Fig. 1A; Table S1). All the pentatomid and scutellerid species are associated with obligatory gut symbiotic bacteria, which are allied to such Enterobacteriaceae genera as *Pantoea*, *Erwinia* and *Enterobacter* (Fig. 1B). Notably, the phylogenetic relationship of the symbiotic bacteria is incongruent with the systematics of the host stinkbugs, reflecting recurrent symbiont acquisitions and replacements in the evolutionary course of the pentatomid and scutellerid stinkbugs (29). Distinct from the pentatomid and scutellerid stinkbugs that vertically transmit their beneficial gut symbiont via egg surface contamination as mentioned above (27, 28), the alydid stinkbug *R. pedestris* does not vertically transmit but environmentally acquires its beneficial Δ-proteobacterial gut symbiont *Caballeronia* (formerly called *Burkholderia*) (18, 30) (Fig. 1B). In this study, for each of these stinkbug species, we generated symbiont-free newborn nymphs and orally inoculated the cultured bacteria to them as depicted in Fig. S1. We collected eggs and surface-sterilized them with ethanol and formalin, from which symbiont-free newborn nymphs hatched. The nymphs were kept without water for 1 day and then provided with bacteria-suspended water for 1 day, by which the nymphs actively ingested the bacteria-suspended water. Then, the nymphs were provided with sterilized water and food seeds in clean rearing containers, and maintained and inspected every day for monitoring survival and adult emergence as shown in Fig. S1 and Table S1. Bacterial infection was surveyed at the 2nd instar stage and the adult stage to confirm the initial infection rate and the final establishment rate, respectively.

**FIG 1.**
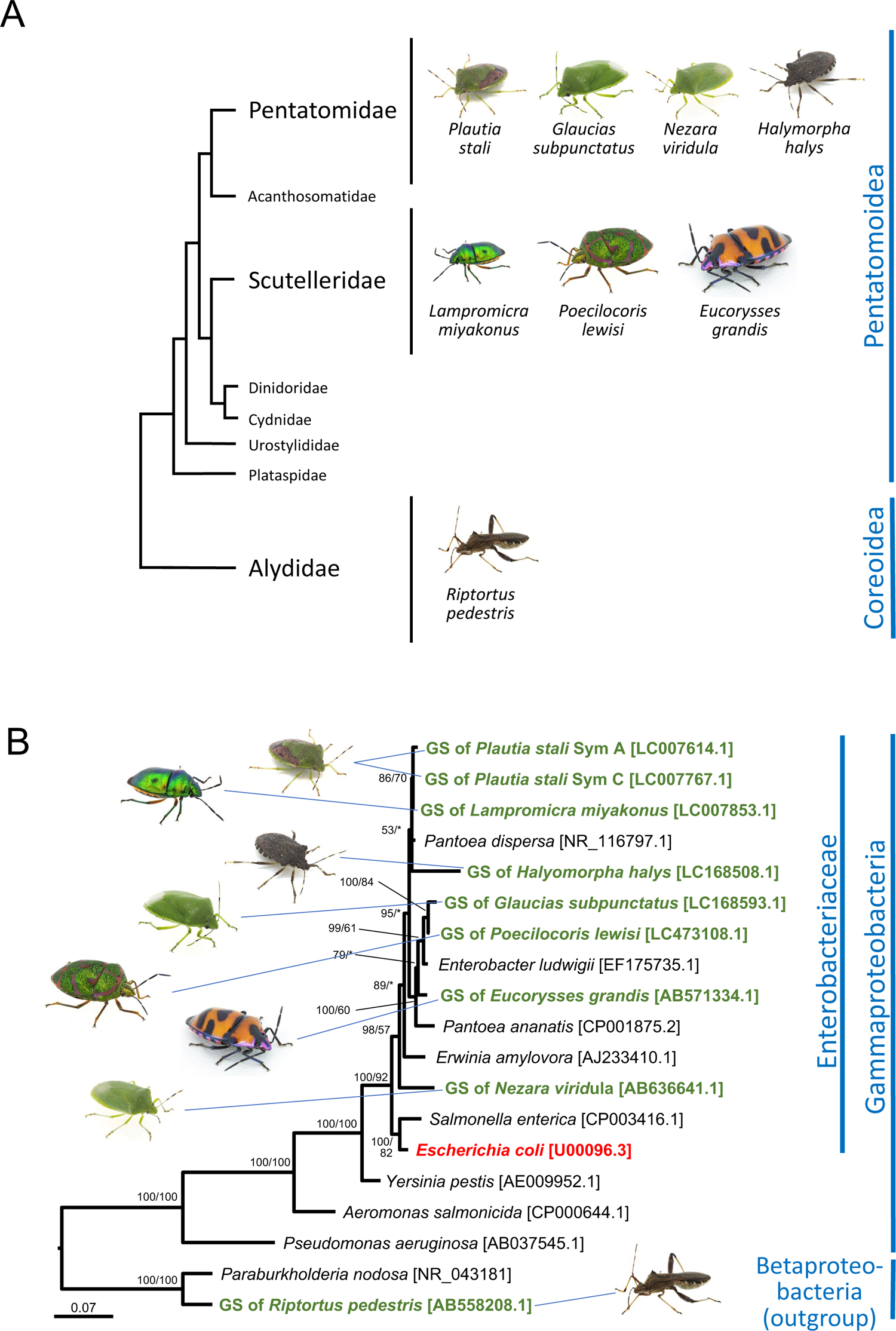
Stinkbugs and their symbiotic bacteria. (**A**) Systematic relationship of the stinkbugs used in this study. The relationship of the stinkbug families is after Wu et al. (45). Stinkbug superfamilies are displayed on the right side. (**B**) Phylogenetic relationship of the symbiotic bacteria. A maximum-likelihood phylogeny inferred from bacterial 16S rRNA gene sequences (1,462 aligned nucleotide sites) is shown. On each node, statistical support values are indicated as posterior probability of Baysian inference/bootstrap probability of maximum-likelihood analysis. Gut symbiotic bacteria (= GS) of the stinkbugs are shown in green, *E. coli* is highlighted in red, and free-living proteobacterial 16S rRNA gene sequences retrieved from the DNA databases are displayed in black. Bacterial families and classes are shown on the right size. Corresponding host stinkbugs are indicated by line connections. Note that the symbiont phylogeny does not agree with the host systematics, reflecting recurrent symbiont acquisitions and replacements in the evolutionary course of the stinkbugs (29).

### Symbiont replacing experiments: *Plautia stali* (Fig. 2; Fig. S2)

The control insects infected with the original symbiont attained high survival rate (88.6%) and adult emergence rate (87.7%). The aposymbiotic insects exhibited high survival rate (83.2%), but actually their growth was very poor: most of them remained at second or third instar with no adult emergence (0%). These results confirmed the importance of the original symbiont Sym A for *P. stali* as reported previously (24). The mutualistic *E. coli* Λ′*cyaA*-inoculated insects showed good survival rate (68.1%) and adult emergence rate (54.9%). The natural cultivable symbiont Sym C-inoculated insects exhibited moderate survival rate (58.0%) and adult emergence rate (28.6%). The wild-type *E. coli* Λ′*intS*-inoculated insects showed moderate survival rate (37.9%) and low adult emergence rate (12.1%). While Λ′*intS E. coli*, Λ′*cyaA E. coli* and Sym C symbiont exhibited moderate PCR detection rates at the second instar stage (37.5%, 54.2% and 37.5%, respectively), they consistently showed 100% detection rates at the adult stage.

**FIG 2.**
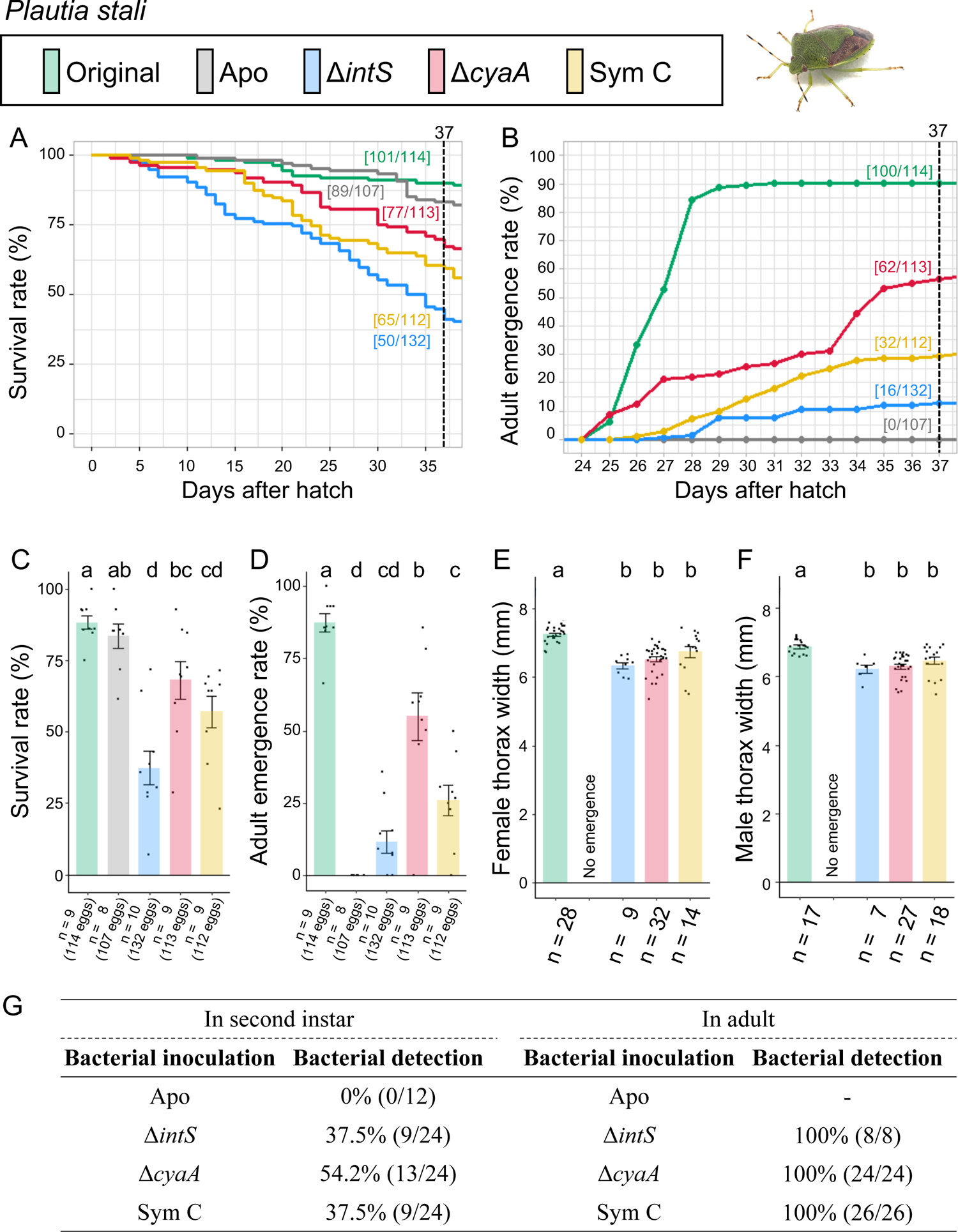
Infection, survival and growth of *P. stali* experimentally deprived of the original mutualistic symbiont and inoculated with either a wild-type *E. coli* strain (Δ*intS*), a mutant *E. coli* strain mutualistic to *P. stali* (Δ*cyaA*), or a natural cultivable symbiont strain mutualistic to *P. stali* (SymC). (**A**) Survival curve. (**B**) Adult emergence curve. (**C**) Survival rate on the 37^th^ day after hatch. (**D**) Adult emergence rate on the 37^th^ day after hatch. (**E**) Adult female body size. (**F**) Adult male body size. (**G**) PCR check of bacterial infection in second instar nymphs (left) and adult insects (right). Statistical tests were conducted by likelihood ratio test of a generalized linear model in (**C**) and (**D**) (a-d, *P* < 0.05) and by pair-wise *t* test with Bonferroni correction in (**E**) and (**F**) (a-b, *P* < 0.05).

These results indicated that (i) both the natural cultivable symbiont Sym C and the mutualistic *E. coli* strain Λ′*cyaA* can establish infection in *P. stali,* and (ii) both Sym C and Λ′*cyaA* can support growth and survival of *P. stali* to adulthood, as reported in previous studies (24, 26).

### Symbiont replacing experiments: *Glaucias subpunctatus* (Fig. 3; Fig. S3)

The control insects infected with the original symbiont attained good survival rate (67.9%) and adult emergence rate (67.9%). The aposymbiotic insects exhibited low survival rate (27.6%) and very low adult emergence rate (3.1%). These results indicated the importance of the original symbiont for *G. subpunctatus*. The mutualistic *E. coli* Λ′*cyaA*-inoculated insects showed good survival rate (64.4%), but adult emergence rate was very low (8.9%). The wild-type *E. coli* Λ′*intS*-inoculated insects similarly showed good survival rate (59.7%) and very low adult emergence rate (0.6%). The Sym C-inoculated insects exhibited relatively low survival rate (21.9%), but adult emergence rate was not so low (18.1%) in comparison with the *E. coli*-infected insects. When inoculated with Λ′*intS* and Λ′*cyaA E. coli* strains, none of second instar nymphs and adult insects retained bacterial infection. On the other hand, Sym C-inoculated insects exhibited relatively low infection rate (16.7%) at the second instar stage and high infection rate (84.6%) at the adult stage.

**FIG 3.**
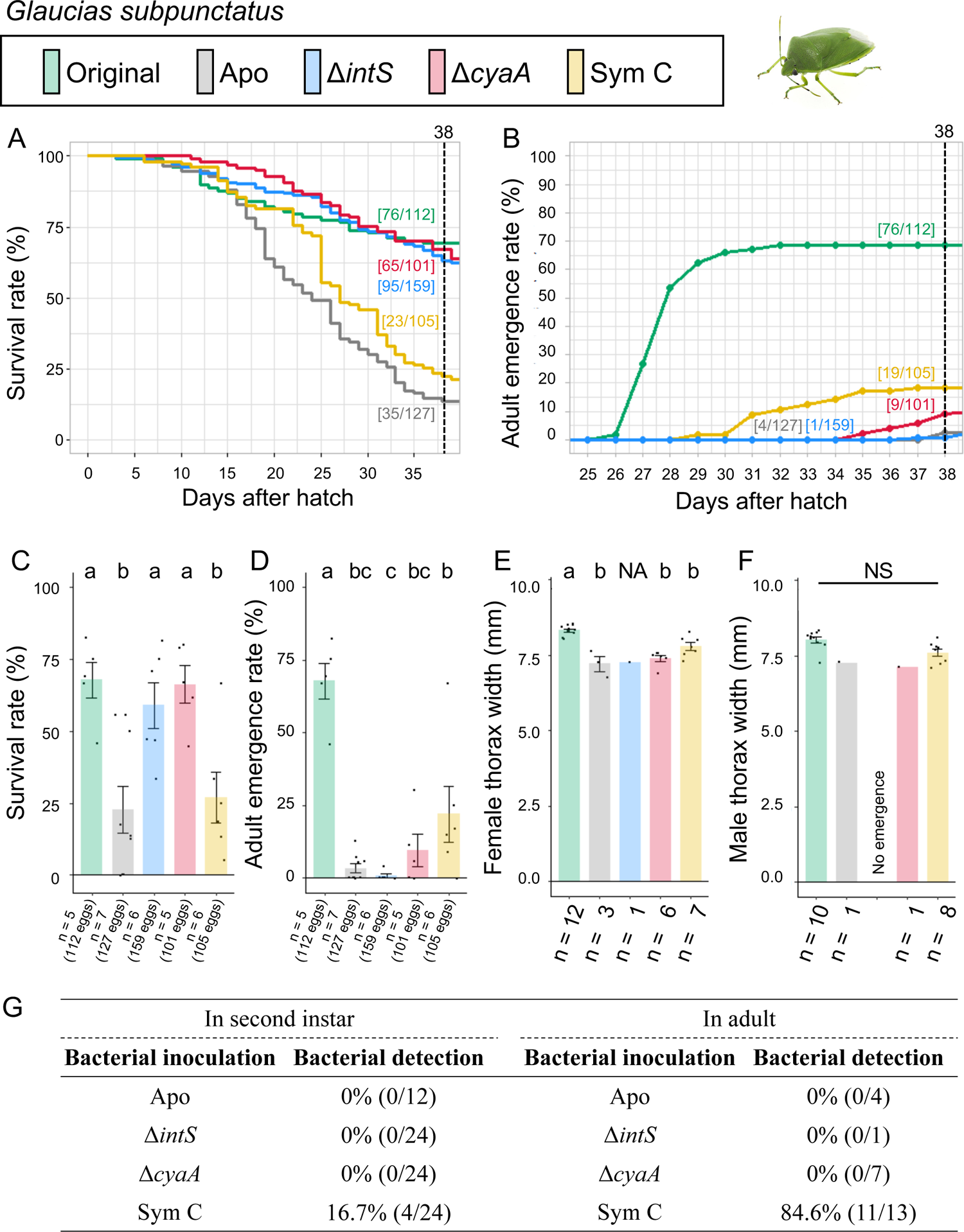
Infection, survival and growth of *G. subpunctatus* experimentally deprived of the original mutualistic symbiont and inoculated with either a wild-type *E. coli* strain (Δ*intS*), a mutant *E. coli* strain mutualistic to *P. stali* (Δ*cyaA*), or a natural cultivable symbiont strain mutualistic to *P. stali* (SymC). (**A**) Survival curve. (**B**) Adult emergence curve. (**C**) Survival rate on the 38^th^ day after hatch. (**D**) Adult emergence rate on the 38^th^ day after hatch. (**E**) Adult female body size. (**F**) Adult male body size. (**G**) PCR check of bacterial infection in second instar nymphs (left) and adult insects (right). Statistical tests were conducted by likelihood ratio test of a generalized linear model in (**C**) and (**D**) (a-c, *P* < 0.05) and by pair-wise *t* test with Bonferroni correction in (**E**) and (**F**) (a-b, *P* < 0.05; NS, no significant difference).

These results indicated that (i) the natural cultivable symbiont Sym C can partially establish infection and support growth and survival of *G. subpunctatus*, and by contrast, (ii) neither the mutualistic *E. coli* Δ*cyaA* nor the wild-type *E. coli* Δ*intS* can establish infection and support growth and survival of *G. subpunctatus*.

### Symbiont replacing experiments: *Nezara viridula* (Fig. 4; Fig. S4)

The control insects infected with the original symbiont exhibited relatively low survival rate (22.6%) and adult emergence rate (21.3%). The relatively low level of performance of the insects infected with the original symbiont suggested that the laboratory rearing system does not work perfectly for this stinkbug species. Notably, however, the aposymbiotic insects ended up with no survivors (0%) and no adult insects (0%), confirming the importance of the original symbiont for *N. viridula* as previously reported (31). The mutualistic *E. coli* Δ*cyaA*-inoculated insects suffered very low survival (6.6%) and few adult emergence (0.8%). The wild-type *E. coli* Δ*intS*-inoculated insects similarly showed very low survival (8.7%) and no adult emergence (0%). The Sym C-inoculated insects exhibited relatively low survival rate (33.1%) and few adult emergence (0.7%). When inoculated with Δ*intS* and Δ*cyaA E. coli* strains, none of second instar nymphs and an adult insect retained bacterial infection. On the other hand, Sym C-inoculated insects exhibited relatively low infection rate (17.4%) at the second instar stage, and the only adult insect obtained was infected with Sym C.

**FIG 4.**
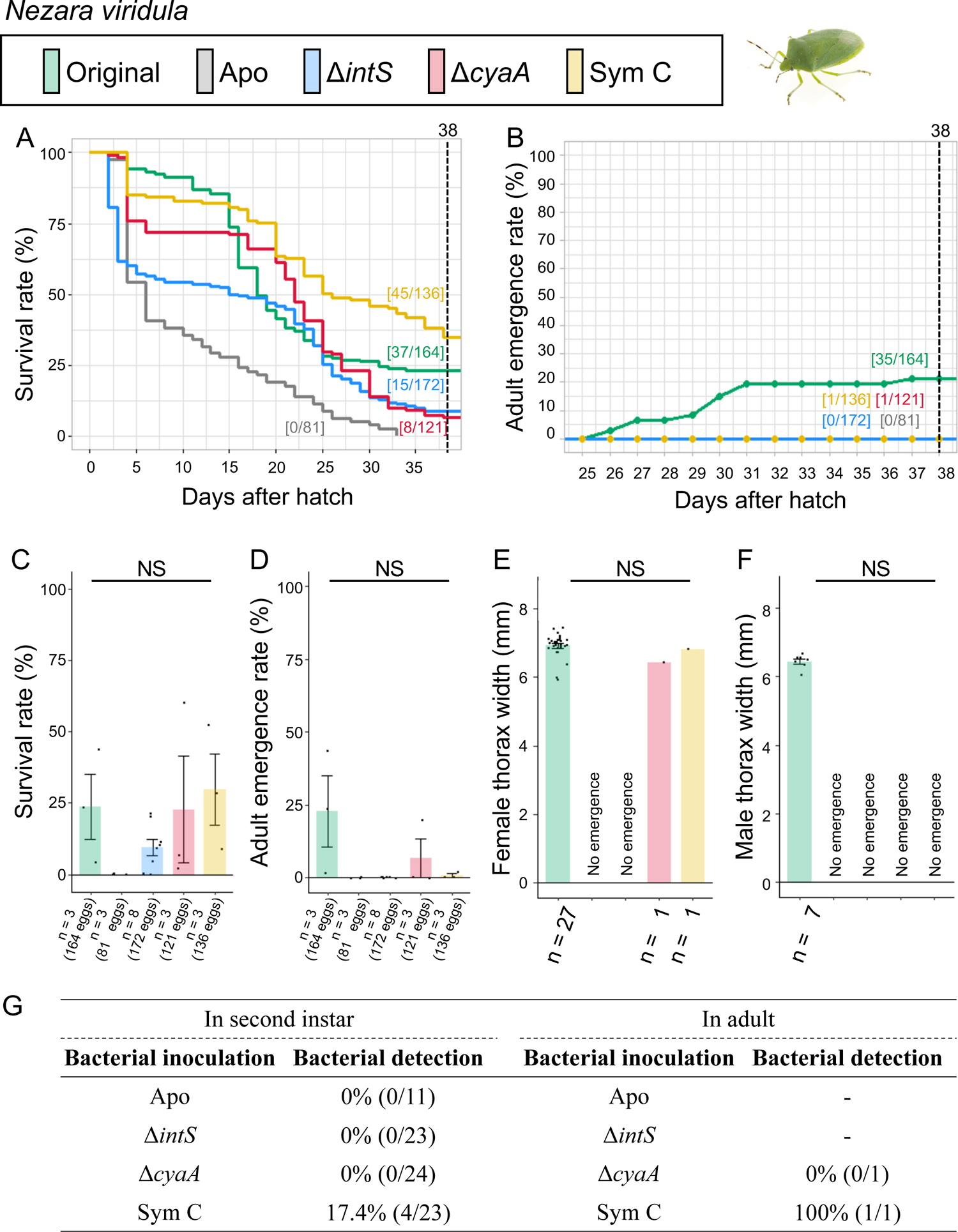
Infection, survival and growth of *N. viridula* experimentally deprived of the original mutualistic symbiont and inoculated with either a wild-type *E. coli* strain (Δ*intS*), a mutant *E. coli* strain mutualistic to *P. stali* (Δ*cyaA*), or a natural cultivable symbiont strain mutualistic to *P. stali* (SymC). (**A**) Survival curve. (**B**) Adult emergence curve. (**C**) Survival rate on the 38^th^ day after hatch. (**D**) Adult emergence rate on the 38^th^ day after hatch. (**E**) Adult female body size. (**F**) Adult male body size. (**G**) PCR check of bacterial infection in second instar nymphs (left) and adult insects (right). Statistical tests were conducted by likelihood ratio test of a generalized linear model in (**C**) and (**D**) (NS, no significant difference) and by pair-wise *t* test with Bonferroni correction in (**E**) and (**F**) (NS, no significant difference).

These results indicated that (i) the natural cultivable symbiont Sym C can establish infection and support survival of *N. viridula* to some extent, and by contrast, (ii) neither the mutualistic *E. coli* Δ*cyaA* nor the wild-type *E. coli* Δ*intS* can establish infection and support growth and survival of *N. viridula*.

### Symbiont replacing experiments: *Halyomorpha halys* (Fig. 5; Fig. S5)

The control insects infected with the original symbiont attained good survival rate (57.1%) and adult emergence rate (38.1%), reflecting the importance of the original symbiont for *H. halys* as previously reported (32). Strikingly, however, although similar survival rates were observed with the aposymbiotic insects (51.0%), the mutualistic *E. coli* Δ*cyaA*-inoculated insects (65.6%), the wild-type *E. coli* Δ*intS*-inoculated insects (64.7%), and the Sym C-inoculated insects (54.2%), none of them attained adulthood. While half of the Δ*cyaA*-inoculated insects retained bacterial infection at the second instar stage, few second instar insects exhibited bacterial infection when inoculated with Δ*intS* or Sym C.

**FIG 5.**
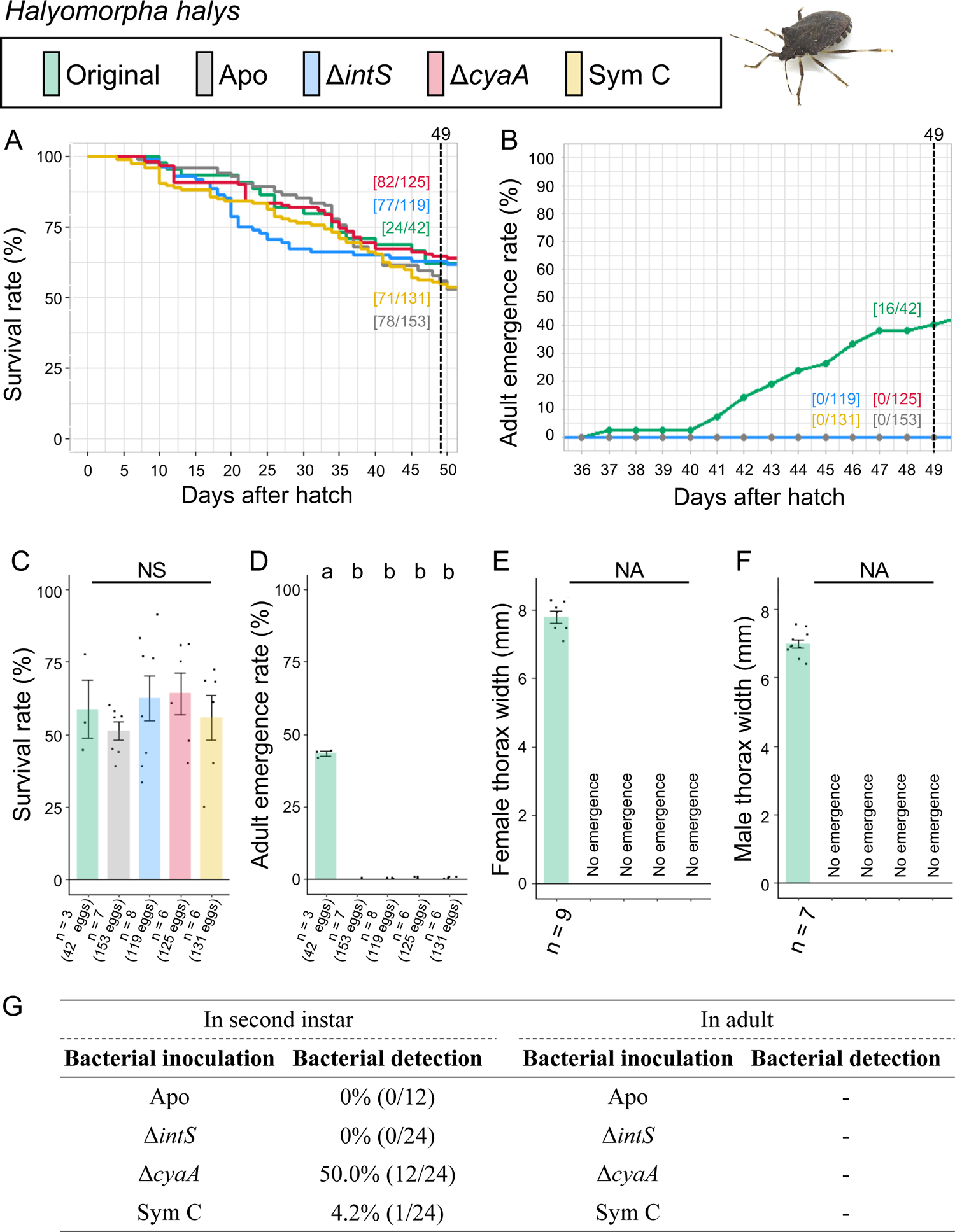
Infection, survival and growth of *H. halys* experimentally deprived of the original mutualistic symbiont and inoculated with either a wild-type *E. coli* strain (Δ*intS*), a mutant *E. coli* strain mutualistic to *P. stali* (Δ*cyaA*), or a natural cultivable symbiont strain mutualistic to *P. stali* (SymC). (**A**) Survival curve. (**B**) Adult emergence curve. (**C**) Survival rate on the 49^th^ day after hatch. (**D**) Adult emergence rate on the 49^th^ day after hatch. (**E**) Adult female body size. (**F**) Adult male body size. (**G**) PCR check of bacterial infection in second instar nymphs (left) and adult insects (right). Statistical tests were conducted by likelihood ratio test of a generalized linear model in (**C**) and (**D**) (a-b, *P* < 0.05; NS, no significant difference). and by pair-wise *t* test with Bonferroni correction in (**E**) and (**F**) (NA, not applicable due to insufficient replicates).

These results indicated that (i) neither the natural cultivable symbiont Sym C, the mutualistic *E. coli* strain Δ*cyaA,* nor the wild-type *E. coli* strain Δ*intS* can establish infection and support growth and survival of *H. halys*, and (ii) in the absence of the original symbiont, *H. halys* can manage to survive but cannot attain adulthood.

### Symbiont replacing experiments: *Lampromicra miyakonus* (Fig. 6; Fig. S6)

The control insects infected with the original symbiont attained good survival rate (55.6%) and adult emergence rate (55.6%). The aposymbiotic insects exhibited low survival rate (1.6%) with no adult emergence (0%). These results confirmed the importance of the original symbiont for *L. miyakonus* as previously reported (33). The natural cultivable symbiont Sym C-inoculated insects efficiently established infection (93.3% in nymphs and 100% in adults) and exhibited moderate survival rate (37.5%) and adult emergence rate (35.9%). By contrast, when inoculated with the mutualistic *E. coli* strain Δ*cyaA* and the wild-type *E. coli* strain Δ*intS*, infection rates at the second instar were partial (20.0% and 46.7%, respectively) and few insects survived and attained adulthood.

**FIG 6.**
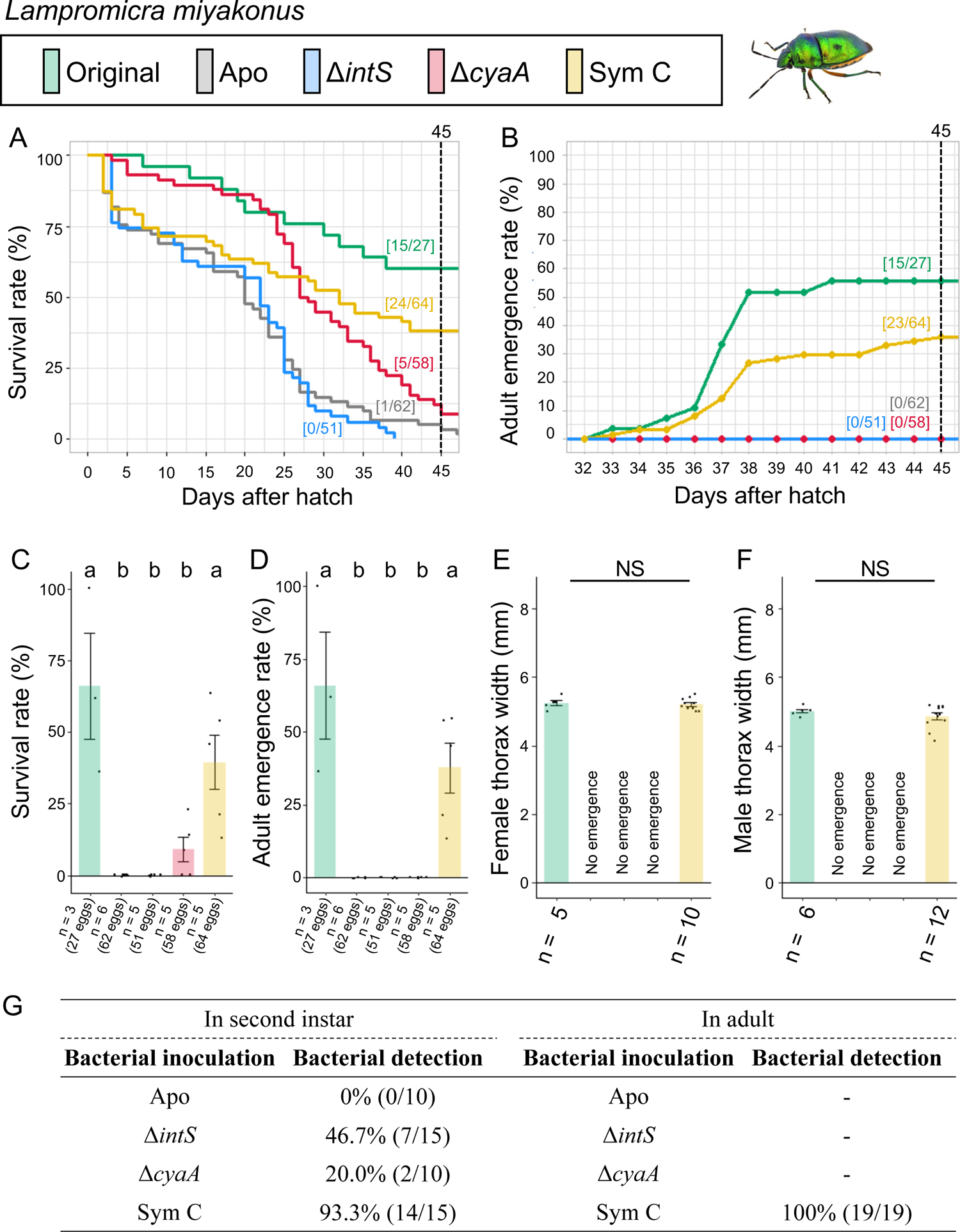
Infection, survival and growth of *L. miyakonus* experimentally deprived of the original mutualistic symbiont and inoculated with either a wild-type *E. coli* strain (Δ*intS*), a mutant *E. coli* strain mutualistic to *P. stali* (Δ*cyaA*), or a natural cultivable symbiont strain mutualistic to *P. stali* (SymC). (**A**) Survival curve. (**B**) Adult emergence curve. (**C**) Survival rate on the 45^th^ day after hatch. (**D**) Adult emergence rate on the 45^th^ day after hatch. (**E**) Adult female body size. (**F**) Adult male body size. (**G**) PCR check of bacterial infection in second instar nymphs (left) and adult insects (right). Statistical tests were conducted by likelihood ratio test of a generalized linear model in (**C**) and (**D**) (a-b, *P* < 0.05) and by pair-wise *t* test with Bonferroni correction in (**E**) and (**F**) (NS, no significant difference).

These results indicated that (i) the natural cultivable symbiont Sym C can establish infection and support survival of *L. miyakonus*, and (ii) neither the mutualistic *E. coli* Δ*cyaA* nor the wild-type *E. coli* Δ*intS* can establish infection and support growth and survival of *L. miyakonus*.

### Symbiont replacing experiments: *Poecilocoris lewisi* (Fig. 7; Fig. S7)

The control insects infected with the original symbiont attained high survival rate (77.8%) and adult emergence rate (77.8%). The aposymbiotic insects exhibited low survival rate (22.2%) and no adult emergence (0%). These results indicated the importance of the original symbiont for *P. lewisi*. The natural cultivable symbiont Sym C-inoculated insects efficiently established infection (91.7% in nymphs and 100% in adults) and exhibited high survival rate (90.5%) and adult emergence rate (90.5%). By contrast, when inoculated with the mutualistic *E. coli* strain Δ*cyaA* and the wild-type *E. coli* strain Δ*intS*, no infection was established and no adult insect emerged.

**FIG 7.**
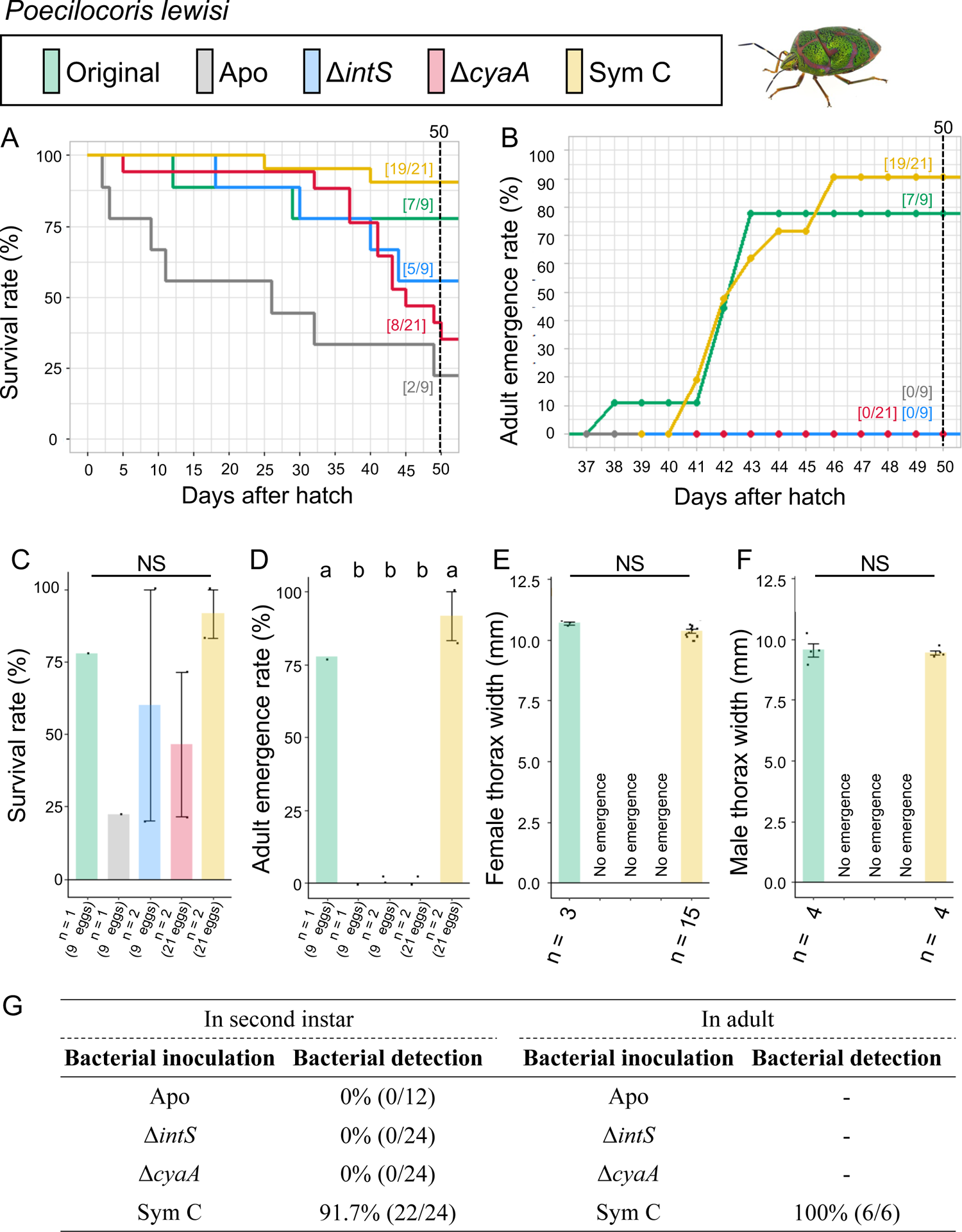
Infection, survival and growth of *P. lewisi* experimentally deprived of the original mutualistic symbiont and inoculated with either a wild-type *E. coli* strain (Δ*intS*), a mutant *E. coli* strain mutualistic to *P. stali* (Δ*cyaA*), or a natural cultivable symbiont strain mutualistic to *P. stali* (SymC). (**A**) Survival curve. (**B**) Adult emergence curve. (**C**) Survival rate on the 50^th^ day after hatch. (**D**) Adult emergence rate on the 50^th^ day after hatch. (**E**) Adult female body size. (**F**) Adult male body size. (**G**) PCR check of bacterial infection in second instar nymphs (left) and adult insects (right). Statistical tests were conducted by likelihood ratio test of a generalized linear model in (**C**) and (**D**) (a-b, *P* < 0.05; NS, no significant difference) and by pair-wise *t* test with Bonferroni correction in (**E**) and (**F**) (NS, no significant difference).

These results indicated that (i) the natural cultivable symbiont Sym C can establish infection and support survival of *P. lewisi*, and (ii) neither the mutualistic *E. coli* Δ*cyaA* nor the wild-type *E. coli* Δ*intS* can establish infection and support growth and survival of *P. lewisi*.

### Symbiont replacing experiments: *Eucorysses grandis* (Fig. 8; Fig. S8)

The control insects infected with the original symbiont attained high survival rate (100%) and adult emergence rate (92.3%). The aposymbiotic insects exhibited moderate survival rate (31.9%) with no adult emergence (0%). These results indicated the importance of the original symbiont for *E. grandis*. By contrast, when inoculated with the natural cultivable symbiont Sym C, the mutualistic *E. coli* strain Δ*cyaA*, and the wild-type *E. coli* strain Δ*intS*, no infection was established and no adult insect emerged.

**FIG 8.**
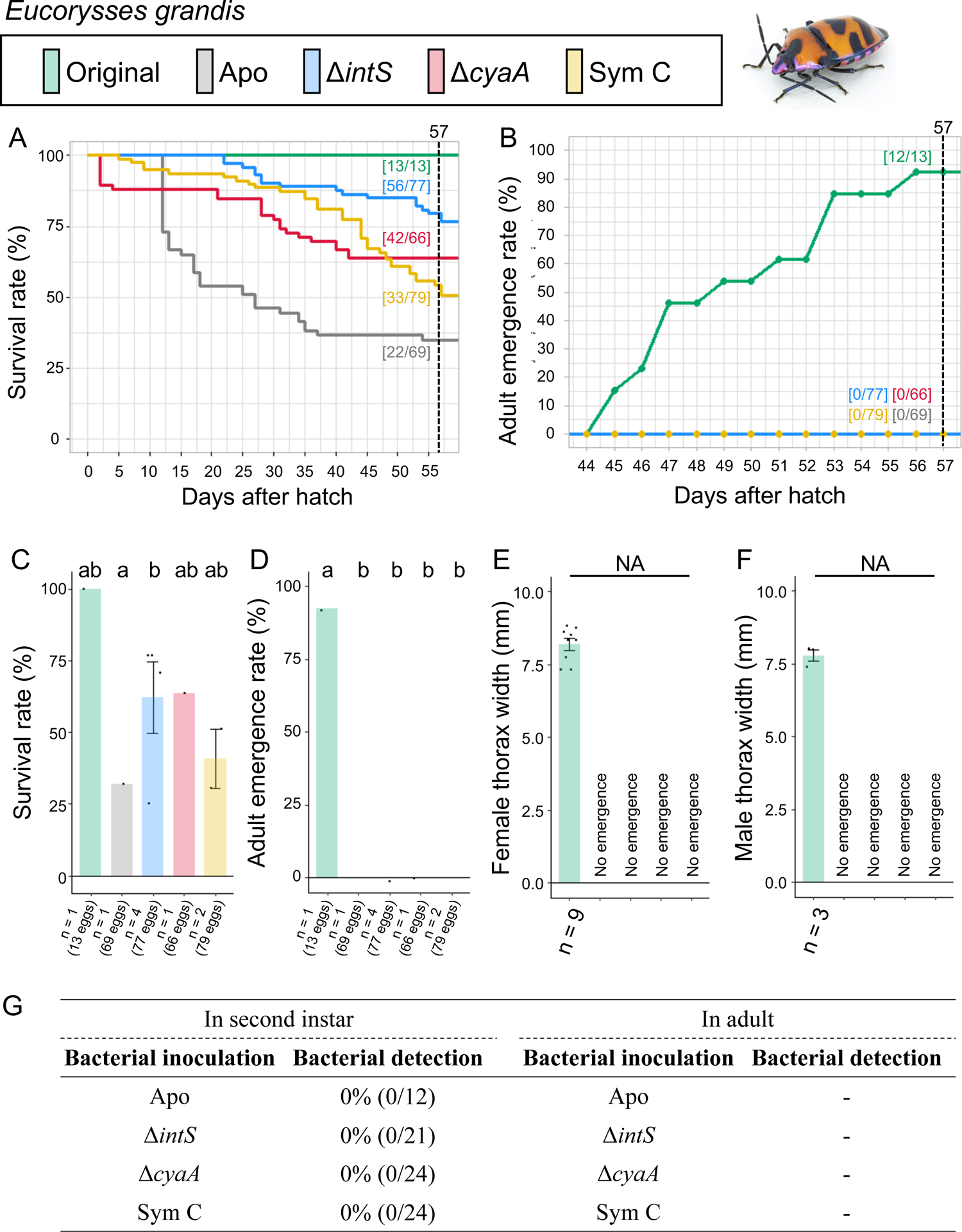
Infection, survival and growth of *E. grandis* experimentally deprived of the original mutualistic symbiont and inoculated with either a wild-type *E. coli* strain (Δ*intS*), a mutant *E. coli* strain mutualistic to *P. stali* (Δ*cyaA*), or a natural cultivable symbiont strain mutualistic to *P. stali* (SymC). (**A**) Survival curve. (**B**) Adult emergence curve. (**C**) Survival rate on the 57^th^ day after hatch. (**D**) Adult emergence rate on the 57^th^ day after hatch. (**E**) Adult female body size. (**F**) Adult male body size. (**G**) PCR check of bacterial infection in second instar nymphs (left) and adult insects (right). Statistical tests were conducted by likelihood ratio test of a generalized linear model in (**C**) and (**D**) (a-b, *P* < 0.05) and by pair-wise *t* test with Bonferroni correction in (**E**) and (**F**) (NA, not applicable due to insufficient replicates).

These results indicated that neither the natural cultivable symbiont Sym C, the mutualistic *E. coli* strain Δ*cyaA,* nor the wild-type *E. coli* strain Δ*intS* can establish infection and support growth and survival of *E. grandis*.

### Symbiont replacing experiments: *Riptortus pedestris* (Fig. 9; Fig. S9)

As for *R. pedestris*, the control insects that emerged from non-sterilized eggs were not examined, because the insects are substantially the same as the aposymbiotic insects derived from surface-sterilized eggs. The aposymbiotic insects exhibited moderate survival rate (54.2%) and adult emergence rate (45.8%). Here it should be noted that aposymbiotic *R. pedestris* can grow to adult and reproduce, although growth rate, body size and fecundity decline in comparison with symbiotic *R. pedestris* (30). When inoculated with the natural cultivable symbiont Sym C, the mutualistic *E. coli* strain Δ*cyaA* and the wild-type *E. coli* strain Δ*intS*, the insects exhibited improved survival rates (69.0%, 72.7% and 63.6%, respectively) and adult emergence rates (62.1%, 72.7% and 57.6%, respectively) in comparison with the aposymbiotic insects, although none of them established infection with the inoculated bacteria.

**FIG 9.**
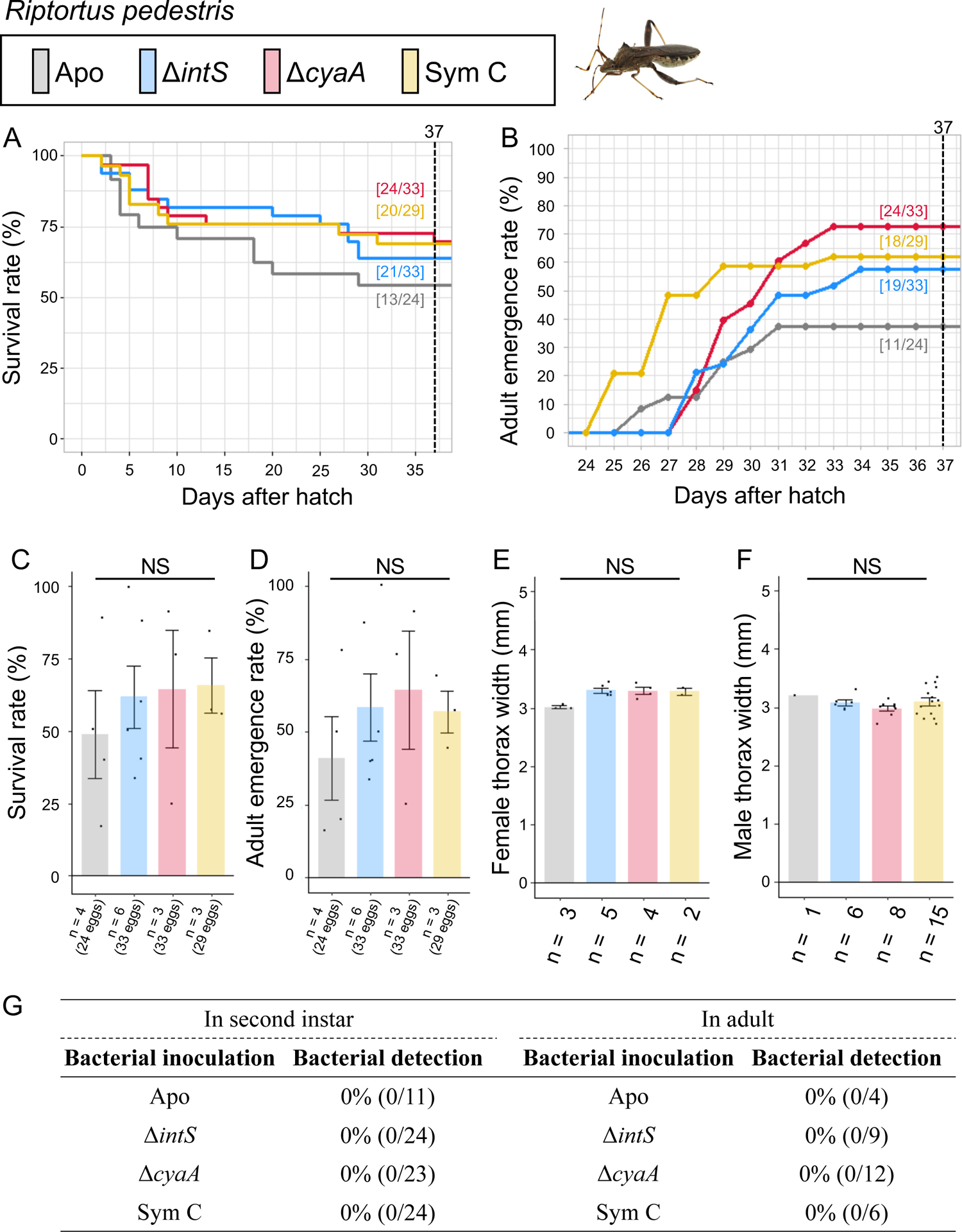
Infection, survival and growth of *E. grandis* experimentally deprived of the original mutualistic symbiont and inoculated with either a wild-type *E. coli* strain (Δ*intS*), a mutant *E. coli* strain mutualistic to *P. stali* (Δ*cyaA*), or a natural cultivable symbiont strain mutualistic to *P. stali* (SymC). (**A**) Survival curve. (**B**) Adult emergence curve. (**C**) Survival rate on the 37^th^ day after hatch. (**D**) Adult emergence rate on the 37^th^ day after hatch. (**E**) Adult female body size. (**F**) Adult male body size. (**G**) PCR check of bacterial infection in second instar nymphs (left) and adult insects (right). Statistical tests were conducted by likelihood ratio test of a generalized linear model in (**C**) and (**D**) (NS, no significant difference) and by pair-wise *t* test with Bonferroni correction in (**E**) and (**F**) (NS, no significant difference).

## DISCUSSION

For understanding the evolution of symbiosis, it is of fundamental interest how host-symbiont specificity has developed during early stages of symbiotic associations. The laboratory model bacterium *E. coli* is originally a mammalian gut microbe and thus unrelated to stinkbugs at all. Hence, the *P. stali-E. coli* experimental symbiotic system we recently established (26) provides an unprecedented opportunity to investigate the early evolutionary stages of symbiosis using the experimentally tractable model insect *P. stali* (34) and the genetically tractable model bacterium *E. coli* (35). In this study, we inoculated the mutualistic *E. coli* strain Δ*cyaA*, which was artificially generated based on the outcome of the *P. stali-E. coli* experimental symbiotic evolution (26), to newborn nymphs of diverse stinkbug species that were experimentally deprived of their original symbiotic bacteria. In addition, we inoculated the natural cultivable symbiont strain Sym C, which is found in southwestern island populations of *P. stali* at low frequencies and likely at a relatively early stage of symbiotic evolution (24), to the symbiont-free newborn nymphs in the same manner. In this way, we were able to experimentally evaluate whether the artificial and natural symbiotic bacteria that have adapted to *P. stali* can establish symbiosis/mutualism with a diverse array of stinkbug species.

Table 1 summarizes the results of the symbiont replacing experiments conducted in this study. The remarkable finding is that *E. coli* can, irrespective of wild-type or mutualistic genotype, establish infection and support growth and survival with *P. stali* only (Fig. 2). All the other stinkbug species we examined are not compatible with *E. coli* symbiotically (Figs. 3-9). This result uncovers that strict host specificity can be established at a very early stage of the evolution of mutualism. Considering our experience that the establishment of the *P. stali-E. coli* experimental symbiotic system was rather straightforward, we planned to launch similar *E. coli* experimental symbiotic systems with other stinkbug species. However, it turned out that such an optimistic idea may not work. It is currently elusive why only *P. stali* can establish infection and symbiosis with *E. coli*. Meanwhile, we now realize that we were lucky to have attempted *E. coli* infection to *P. stali* from the beginning (24) and it serendipitously worked so nicely that the *P. stali-E. coli* experimental symbiotic system was established successfully (26).

**Table 1.** Summary of the results of the symbiont replacing experiments.

| Insect species | $\Delta intS$: Wild-type<br>*E. coli* strain | $\Delta cyaA$: *E. coli*<br>mutant strain<br>mutualistic to *P. stali* | Sym C: Natural<br>cultivable symbiont<br>mutualistic to *P. stali* |
| --- | --- | --- | --- |
| **Pentatomidae** |  |  |  |
| *Plautia stali* | △ | ○ | ○ |
| *Glaucias subpunctatus* | × | × | △ |
| *Nezara viridula* | × | × | × |
| *Halyomorpha halys* | × | × | × |
| **Scutelleridae** |  |  |  |
| *Lampromicra miyakonus* | × | × | ○ |
| *Poecilocoris lewisi* | × | × | ○ |
| *Eucorysses grandis* | × | × | × |
| **Alydidae** |  |  |  |
| *Riptortus pedestris* | × | × | × |

○: Infection establishes, and host growth and survival are supported well.
△: Infection establishes to some extent, and host growth and survival are supported partially.
×: Infection does not establish, and host growth and survival are not supported.

Another remarkable finding is that Sym C, the natural cultivable symbiont of *P. stali,* can establish infection and support growth and survival not only with *P. stali* but also with other stinkbug species (Table 1). In *L. miyakona* and *P. lewisi*, Sym C inoculation significantly improved survival and adult emergence (Figs. 6 and 7), whereas the improvement was only partial in *G. subpunctatus* (Fig. 3). Notably, these results can be explained neither by phylogenetic affinity of the bacterial side nor by phylogenetic affinity of the host side. As for the host side, *L. miyakona* and *P. lewisi* belong to the family Scutelleridae that is distinct from the family Pentatomidae to which *P. stali* belongs (Fig. 1A), but *P. stali*-derived Sym C worked well in *L. miyakona* and *P. lewisi* symbiotically (Figs. 6 and 7). As for the bacterial side, certainly the original symbiont of *L. miyakona* is closely related to Sym C, but the original symbionts of *P. lewisi* and *G. subpunctatus* belong to a different clade in the Enterobacteriaceae (Fig. 1B).

Why is Sym C capable of symbiosis with more diverse stinkbug species than *E. coli*? Although speculative, we point out several candidate factors potentially relevant to the different levels of host specificity. A candidate factor is the phylogenetic one. *E. coli* is outside the *Pantoea*/*Enterobacter*/*Erwinia* clade to which the majority of pentatomid and scutellerid symbionts, including Sym C, belong (Fig. 1B) (29). This may account for the limited symbiotic capability of *E. coli* in comparison with *Pantoea*-applied Sym C. Another candidate factor is the historical/co-evolutionary one. In the southwestern Ryukyu islands, the same cultivable symbiotic bacteria, including Sym C, Sym D and Sym E, are shared among different stinkbug species, including *P. stali*, *L. miyakonus*, *Axiogastus rosmatus* and *Solenosthedium chinense*, and also detected from environmental soil samples (24). This situation suggests the possibility that these cultivable symbiotic bacteria may be ecologically shared among the different stinkbug species via heterospecific horizontal transfers or soil-mediated environmental acquisitions. The broader host insect range of Sym C may make sense in this context.

In this study, to our surprise, the mutualistic *E. coli* strain Δ*cyaA* supported survival and adult emergence of *P. stali* significantly better than Sym C that is the natural cultivable symbiont of *P. stali* (Fig. 2). It seems plausible, although speculative, that this unexpected situation is partly ascribed to the excellent symbiotic capability of the mutualistic *E. coli* strain Δ*cyaA* (26), and partly due to the fact that Sym C is derived from a southwestern island population of *P. stali* and experimentally inoculated to the laboratory strain derived from a mainland population of *P. stali*, where host-symbiont local adaptations may matter (24).

It is notable that, although the effects may be relatively minor, in multiple stinkbug species such as *G. subpunctatus*, *N. viridula*, *P. lewisi*, *E. grandis* and *R. pedestris*, bacterial inoculation was neither established nor contributing to adult emergence, but supportive for survival to some extent (Figs. 3, 4, 7, 8 and 9). These effects may be attributable to some trace nutritional elements like vitamins derived from the bacterial inoculum, which should be experimentally verified in future studies.

In conclusion, a series of symbiont replacing experiments using diverse stinkbug species presented in this study, wherein their original symbiont was replaced by an artificial symbiont or a natural symbiont mutualistic to the specific stinkbug species *P. stali*, shed light on possible evolutionary trajectories toward establishment of host specificity at the onset of symbiosis. The finding that the *E. coli* mutant strain mutualistic to *P. stali* cannot establish symbiosis with other stinkbug species uncovered that host specificity can be established at a very early stage of symbiotic evolution. On the other hand, the finding that the natural symbiont Sym C can establish symbiosis with some of the diverse stinkbug species in addition to *P. stali* highlighted that a broader host range of the symbiont can evolve in nature. These observations based on the experimental and evolutionary manipulations of symbiosis provide invaluable insights into how host-symbiont specificity has been established in the evolutionary course toward elaborate symbiotic associations.

## MATERIALS AND METHODS

### Insect strains

All the insects used in this study, namely *P. stali, G. subpunctatus*, *N. viridula*, *H. halys*, *L. miyakonus*, *E. grandis*, *P. lewisi* and *R. pedestris*, are inbred laboratory strains maintained at least for three years at 25℃ under a long-day regimen (16 h light and 8 h dark) in climate chambers (TOMY, CLE305 or PHCBI, MLR-352) with sterilized water containing 0.05% ascorbic acid (DWA) and food seeds as described (Table S1; Fig. S1).

### Bacterial strains

Considering that the mutualistic *E. coli* lines obtained through evolutionary experiments are under the hyper-mutating Δ*mutS* genetic background (26), it was expected that they would accumulate many mutations and quickly change their genetic and phenotypic properties during the experiments. Therefore, we obtained and used the Δ*cyaA E. coli* mutant strain from the Keio single-gene knockout mutant library (36). This *E. coli* mutant is disruptive of adenylate cyclase (*cyaA*) gene under the wild-type *E. coli* genetic background, which was shown to be mutualistic when inoculated to *P. stali* (26). The Δ*intS E. coli* strain was used as a wild-type control *E. coli* strain. The mutualistic symbiont strain, *Pantoea* sp. C (abbreviated as Sym C), which was closely related to *Pantoea dispersa*, easily cultivable, and isolated from an adult insect of *P. stali* collected at Ishigaki Island, Okinawa, Japan in 2009 (24), was also used as a natural symbiont of *P. stali*. The 16S rRNA gene sequences of these bacteria were obtained from the NCBI database (https://www.ncbi.nlm.nih.gov/) with those of representative proteobacterial species. Phylogenetic relationship of these sequences was constructed using RAxML-ng v1.2.1 (37) and MrBayes v3.2.7a (38) for maximum likelihood analysis and Baysian inference, respectively.

### Generation of symbiont-free newborn nymphs

Egg masses were collected from rearing cases of the insects between 13:00 and 15:00 every day. The pentatomid and scutellerid stinkbugs, *P. stali, G. subpunctatus*, *N. viridula*, *H. halys*, *L. miyakonus*, *E. grandis* and *P. lewisi*, vertically transmit their symbiotic bacteria to offspring by smearing symbiont-containing secretion onto egg surface upon oviposition (27). We sterilized their egg masses by soaking in 70% ethanol for 5-10 sec, treating with 4% formaldehyde for 5 min, and washing in sterilized water for 5 min. After air-dried, each egg mass was kept in a sterilized plastic Petri dish, by which symbiont-free newborn nymphs were obtained. Although the alydid stinkbug *R. pedestris* acquires its bacterial symbiont from surrounding environment every generation (30), we treated the eggs of *R. pedestris* in the same way to obtain symbiont-free newborn nymphs.

### Bacterial inoculation to newborn nymphs

The bacterial inoculation procedures are summarized in Fig. S1. Each bacterial strain, Δ*intS,* Δ*cyaA* or Sym C, was cultured in liquid LB medium at 25°C overnight, and diluted with sterilized water to OD600 = 0.05. In each plastic Petri dish, the symbiont-free newborn nymphs, which hatched from the surface sterilized eggs, were kept for a day without water. Then, a cotton pad was introduced to the Petri dish, to which 0.75 ml of the bacterial suspension was applied, and the nymphs were allowed to orally acquire the bacterial suspension overnight. Next day, the cotton pad for inoculation was removed and 3 ml of sterilized DWA was added to another clean cotton pad, from which the nymphs were allowed to take water. Upon molting to the second instar, some nymphs were subjected to 2^nd^ instar PCR infection check, while other nymphs were transferred to new clean rearing containers and aseptically maintained with sterilized food seeds and DWA as described (Table S1) (34).

### Infection check of second instar nymphs

Second instar nymphs three days after molting were individually subjected to DNA extraction and PCR detection of either Δ*intS*, Δ*cyaA* or Sym C. Each insect was homogenized in a plastic tube with 100 μl of lysis buffer (150 mM NaCl, 10 mM Tris-HCl [pH 8.0], 1 mM EDTA, 0.1% SDS), and subjected to Proteinase K digestion (100 μg/ml) at 56°C overnight. The lysate was extracted with water-saturated phenol-chloroform-isoamylalcohol (25:24:1), precipitated and washed with ethanol, air-dried, and dissolved in 200 μl of DNA suspension buffer (10 mM Tris-HCl [pH 8.0], 0.1 mM EDTA). To detect infection with Δ*intS* or Δ*cyaA*, a 0.22 kb region of Tn5 gene inserted into the *E. coli* genome was amplified by PCR with the primers Tn5-1789F (5’-TGC TCG ACG TTG TCA CTG AA-3’) and Tn5-1879R (5’-GCA GGA GCA AGG TGA GAT GA-3’) (26). PCR was performed with Gflex (TaKaRa) under the temperature profile of an initial denaturation at 94°C for 1 min followed by 30 cycles of 98°C for 10 s, 60°C for 15 s and 68°C for 30 s, and a final extension at 68°C for 5 min. To detect infection with Sym C, a 0.48 kb region of *groEL* gene was amplified by PCR with the primers SymC_F (5’-GAG CTG GAA GAC AAG TTC GAG-3’) and SymC_R (5’-ATG AAT GGG CTT TCC ARC TCC-3’) (24) under the same temperature profile. For quality check of the DNA samples, a 1.6 kb region of insect mitochondrial cytochrome oxidase I (COI) gene was amplified by PCR with the primers mt16SA1 (5’-AAW AAA CTA GGA TTA GAT ACC CTA-3’) and mt16SB1 (5’-TCT TAA TYC AAC ATC GAG GTC GCA A-3’) (39) under the temperature profile of an initial denaturation at 94°C for 1 min followed by 30 cycles of 98°C for 10 s, 55°C for 15 s and 68°C for 2 min, and a final extension at 68°C for 5 min.

### Rearing, morphometry and infection check of adult insects

The nymphs inoculated with either Δ*intS*, Δ*cyaA* or Sym C, together with the untreated control nymphs with the original symbiont and the aposymbiotic control nymphs sterilized without bacterial inoculation, were aseptically maintained with sterilized food seeds and DWA (see Table S1) as described (34). During the maintenance, we recorded the number and the developmental stage of surviving insects from 15:00 to 20:00 every day. All the adult insects were anaesthetized on ice and photographed from the dorsal side using a digital scanner (EPSON, GT-X980). Using the images, their thorax width was measured using the image analyzing software Natsumushi V.1.10 (40). Some of the adult insects were subjected to dissection of their symbiotic organ in PBS. The dissected symbiotic organs were photographed to record color and size, and individually subjected to DNA extraction using QIAamp DNA mini Kit (Qiagen). A 1.5 kb region of bacterial 16S rRNA gene was amplified by PCR with the primers 16SA1 (5′-AGA GTT TGA TCM TGG CTC AG-3′) and 16SB1 (5′-TAC GGY TAC CTT GTT ACG ACT T-3′) (41). PCR was performed with Ex Taq (TaKaRa) under the temperature profile of an initial denaturation at 95°C for 2 min followed by 35 cycles of 95°C for 30 s, 52°C for 1 min and 72°C for 2 min and a final extension at 72°C for 5 min. The PCR products were treated with exonuclease (NEB) and shrimp alkaline phosphatase (TaKaRa), and sequenced using the primers 16SA1 and 16SB1. Sanger sequencing analyses were outsourced (Eurofins Japan). To confirm infection with *E. coli* or Sym C, obtained 16S rRNA gene sequences were compared with the reference sequences using MEGA ver. 11.0 (42).

### Statistical analyses and figure preparations

Statistical analyses were conducted using R ver.4.2.3 (43) and RSudio ver. 2023.06.0+421 (44). Images of insects and isolated midgut specimens were taken on a dissection microscope S9D with FLEXCAM C1 (Leica) controlled with ver. LAS-X (Leica). The obtained images were adjusted manually using Affinity Photo ver. 2.3.0 (Serif Ltd) and arranged using Affinity Designer ver. 2.3.1 (Serif Ltd).

## Supporting information

FiguresS1-S9 and Table S1

## SUPPLEMENTARY MATERIALS

**FIG S1** Insect rearing and bacterial inoculation procedures.

**FIG S2** Adult insects of *P. stali* and their symbiotic organs obtained in this study.

**FIG S3** Adult insects of *G. subpunctatus* and their symbiotic organs obtained in this study.

**FIG S4** Adult insects of *N. viridula* and their symbiotic organs obtained in this study.

**FIG S5** Adult insects of *H. halys* and their symbiotic organs obtained in this study.

**FIG S6** Adult insects of *L. miyakonus* and their symbiotic organs obtained in this study.

**FIG S7** Adult insects of *P. lewisi* and their symbiotic organs obtained in this study.

**FIG S8** Adult insects of *E. grandis* and their symbiotic organs obtained in this study.

**FIG S9** Adult insects of *R. pedestris* and their symbiotic organs obtained in this study.

**TABLE S1** Laboratory strains of stinkbugs used in this study.

## ACKNOWLEDGMENTS

We thank Harumi Yamazaki, Tetsuhiro Hachikawa and Sakiko Toyoda for helping maintenance of stinkbug strains. This study was supported by the Japan Science and Technology Agency ERATO grant no. JPMJER1902 to T.F. and R.K.

