## Supplementary material for "Host range of naturally and artificially evolved symbiotic bacteria for a specific host insect": FiguresS1-S9 and Table S1

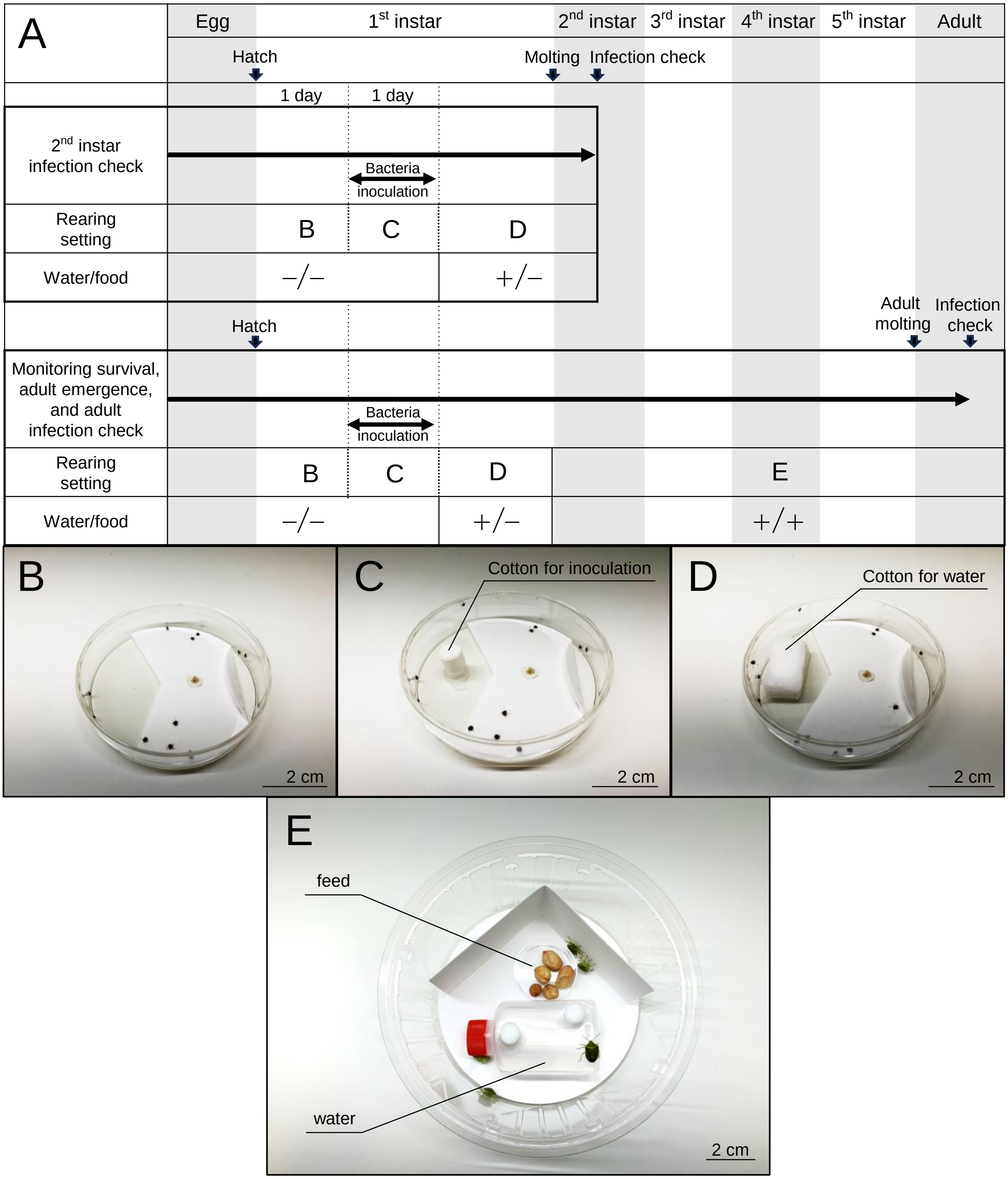


**FIG S1** Insect rearing and bacterial inoculation procedures. (**A**) Procedures of insect rearing, bacterial inoculation, watering and feeding, etc. The upper half shows the procedures for 2nd instar infection check, whereas the bottom half shows the procedures for monitoring survival, adult emergence and adult infection check. (**B-D**) Rearing Petri dish settings for bacterial inoculation to symbiont-free newborn nymphs. (**B**) Before inoculation for 1 day without water. (**C**) During inoculation for 1 day with a cotton pad soaked with bacteria-suspended water. (**D**) After inoculation with a cotton pad soaked with sterile water. (**E**) Rearing container setting for monitoring survival, adult emergence and adult infection check with a water bottle and food seeds.

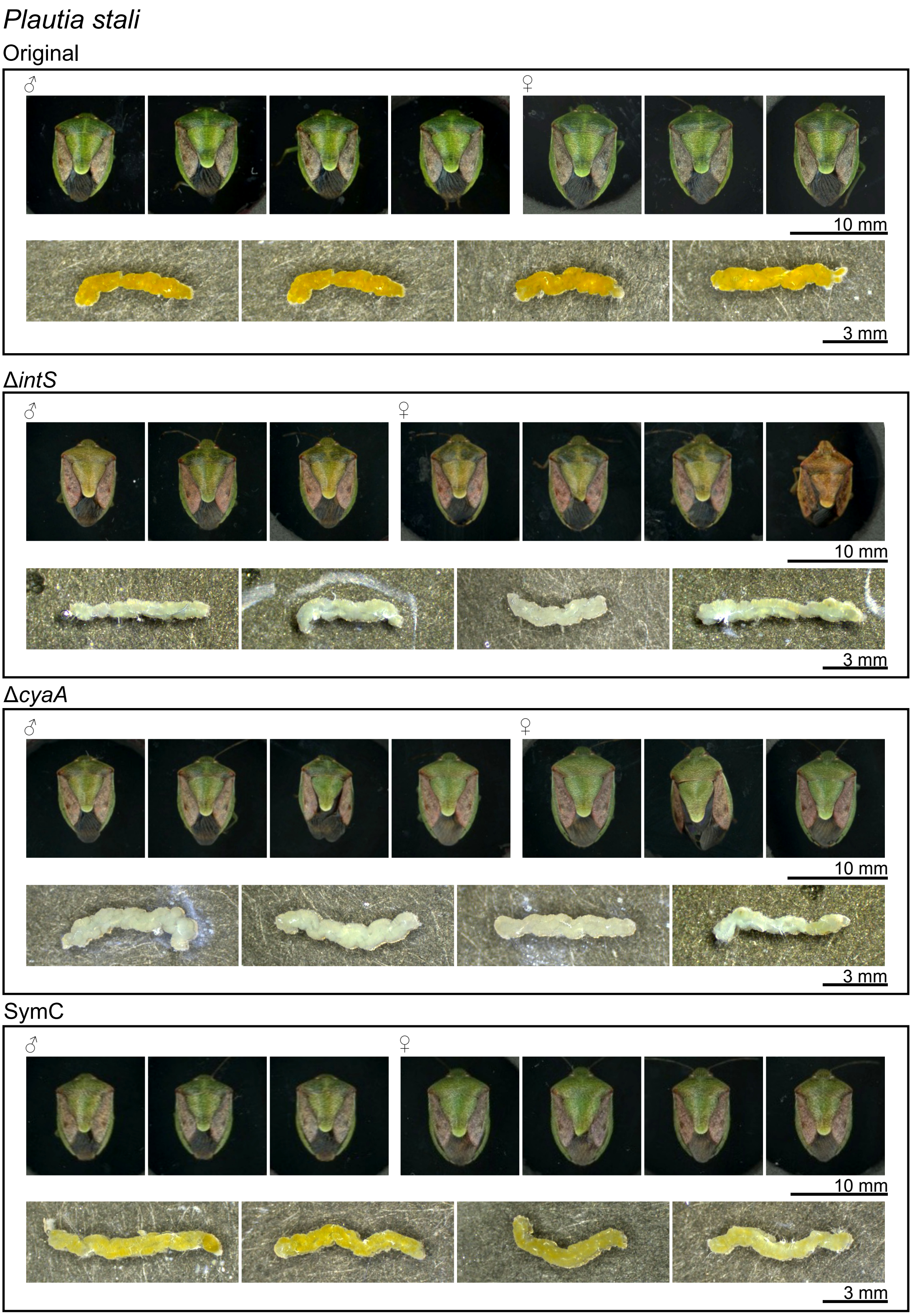


**FIG S2** Adult insects of *P. stali* and their symbiotic organs obtained in this study. Also see Fig. 2.

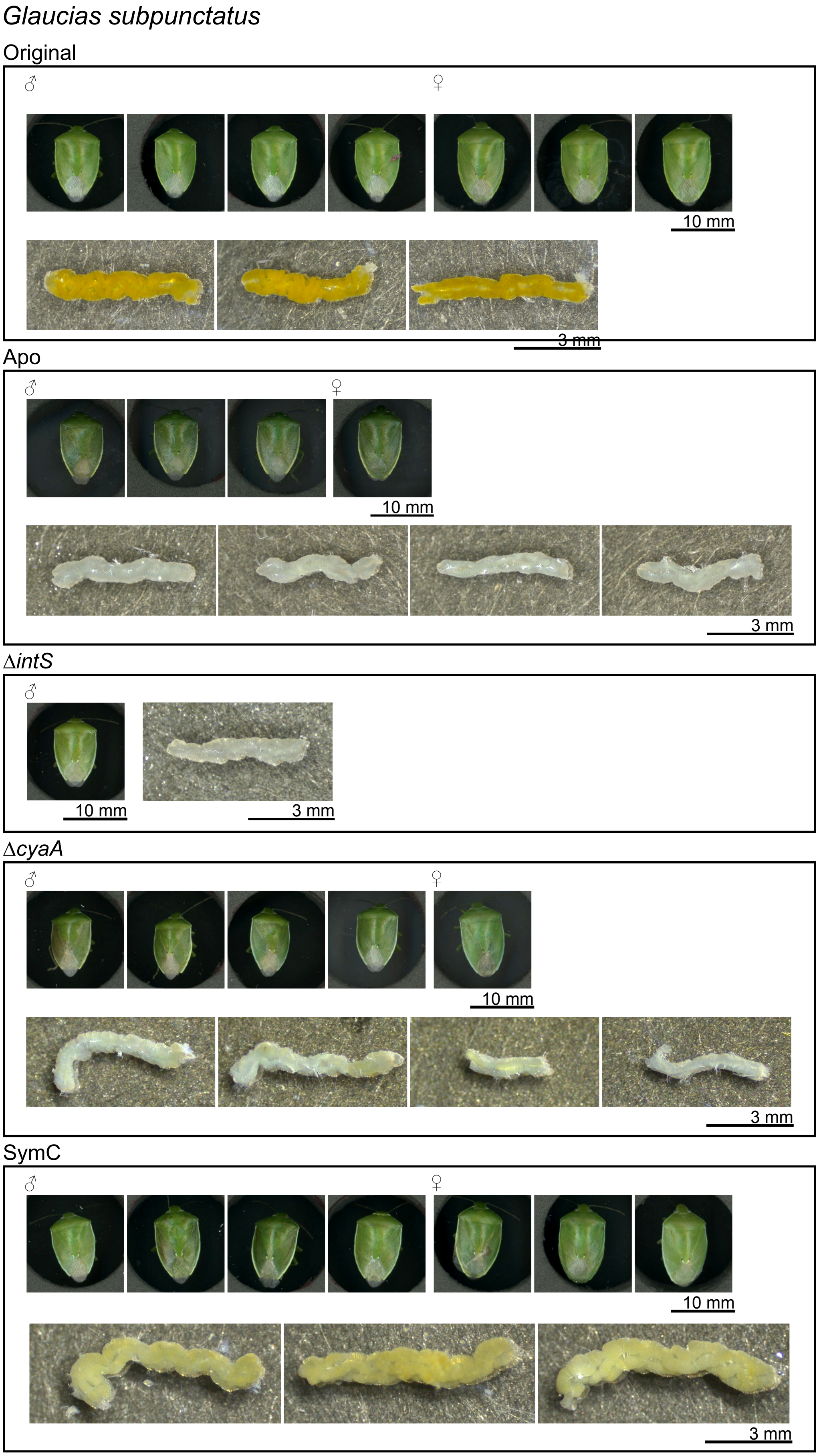


**FIG S3** Adult insects of *G. subpunctatus* and their symbiotic organs obtained in this study. Also see Fig. 3.

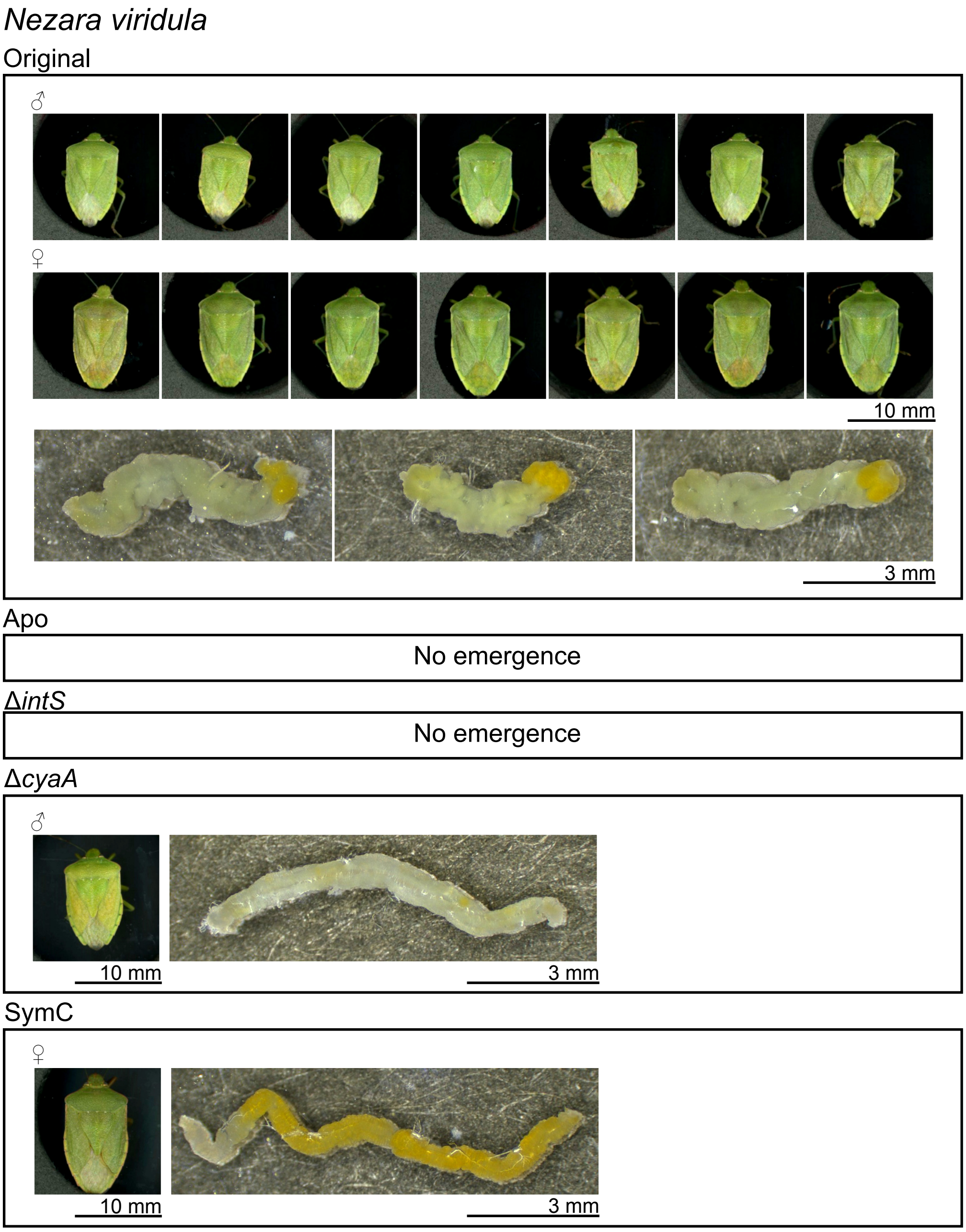


**FIG S4** Adult insects of *N. viridula* and their symbiotic organs obtained in this study. Also see Fig. 4.

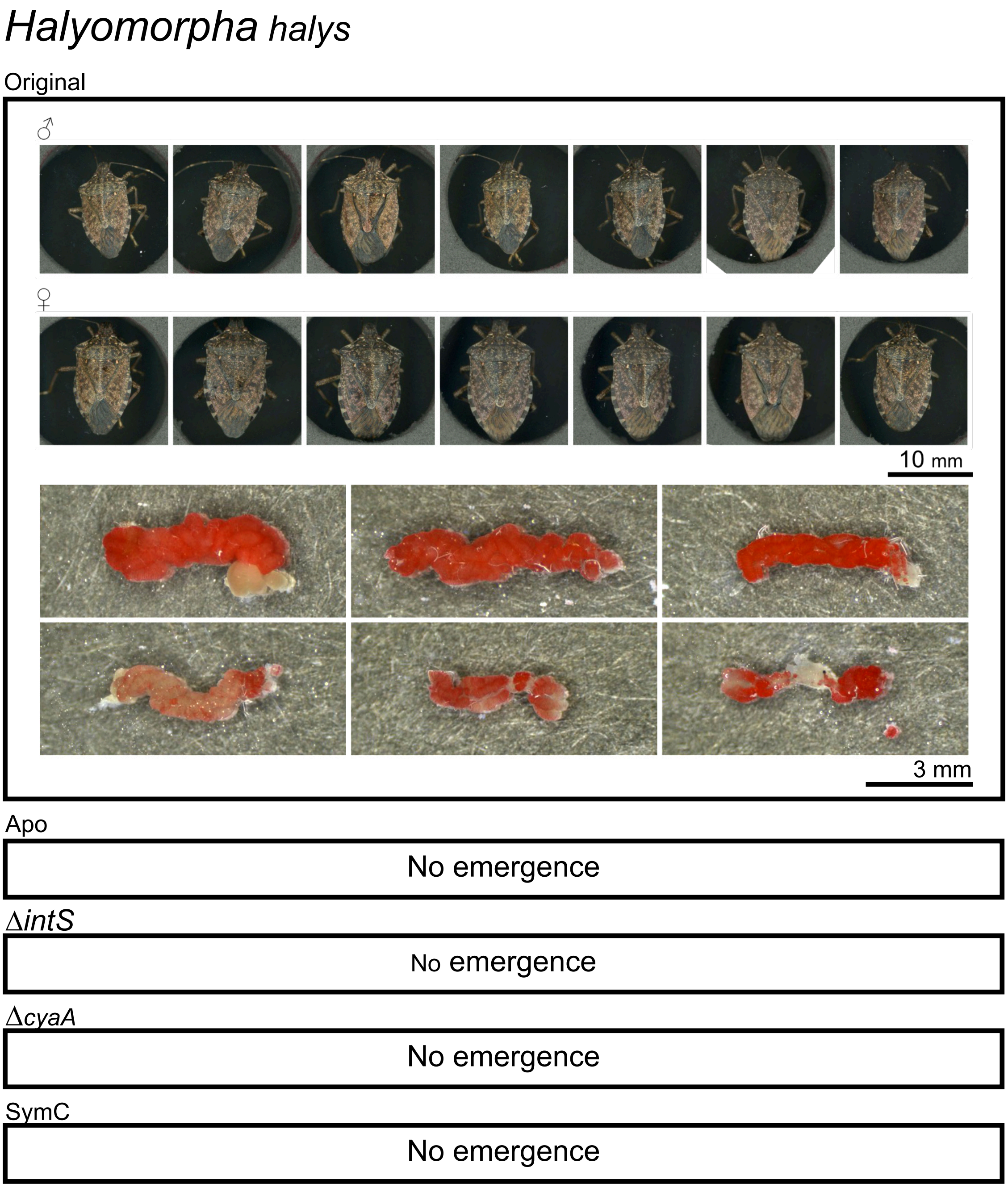


**FIG S5** Adult insects of *H. halys* and their symbiotic organs obtained in this study. Also see Fig. 5.

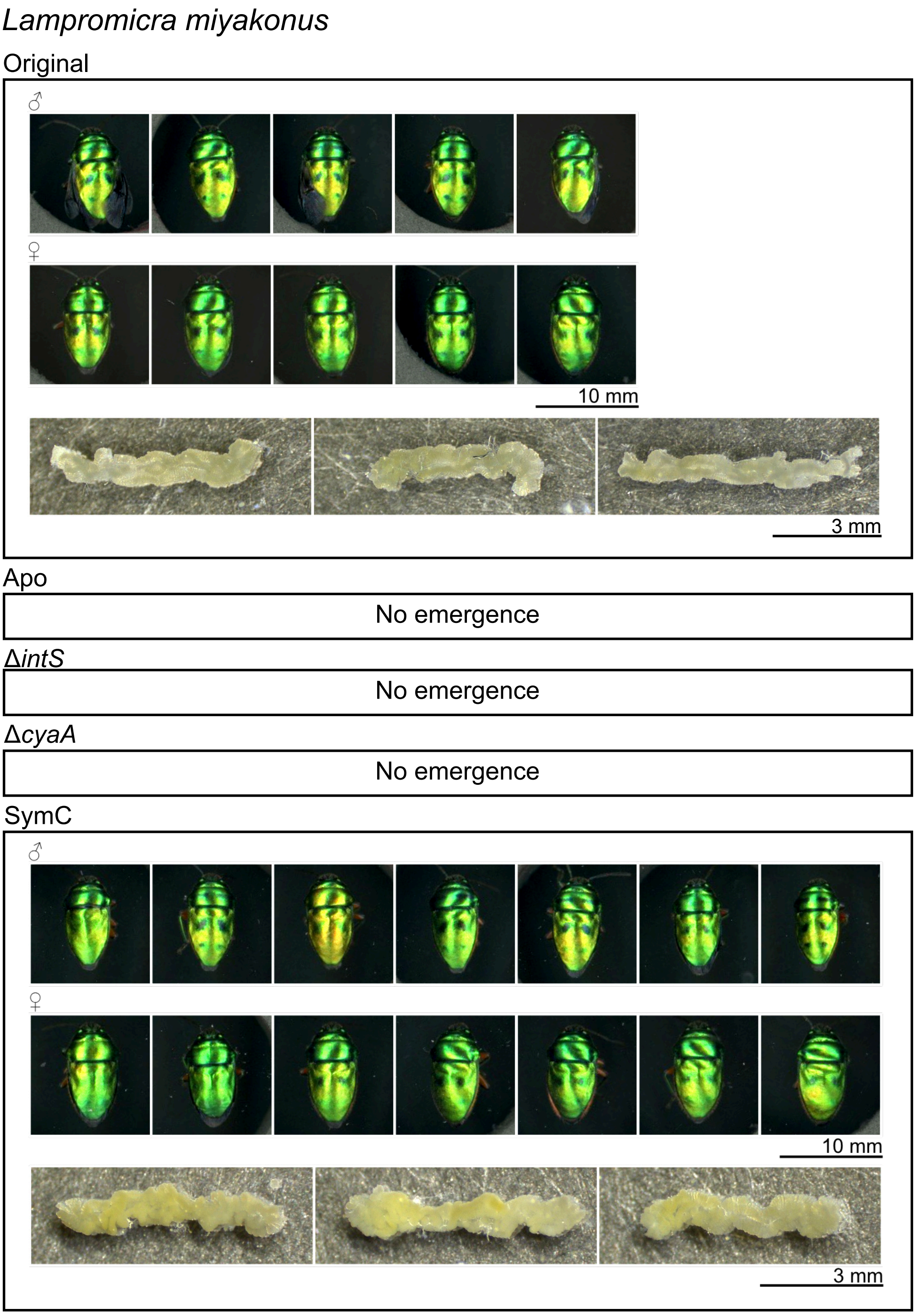


**FIG S6** Adult insects of *L. miyakonus* and their symbiotic organs obtained in this study. Also see Fig. 6.

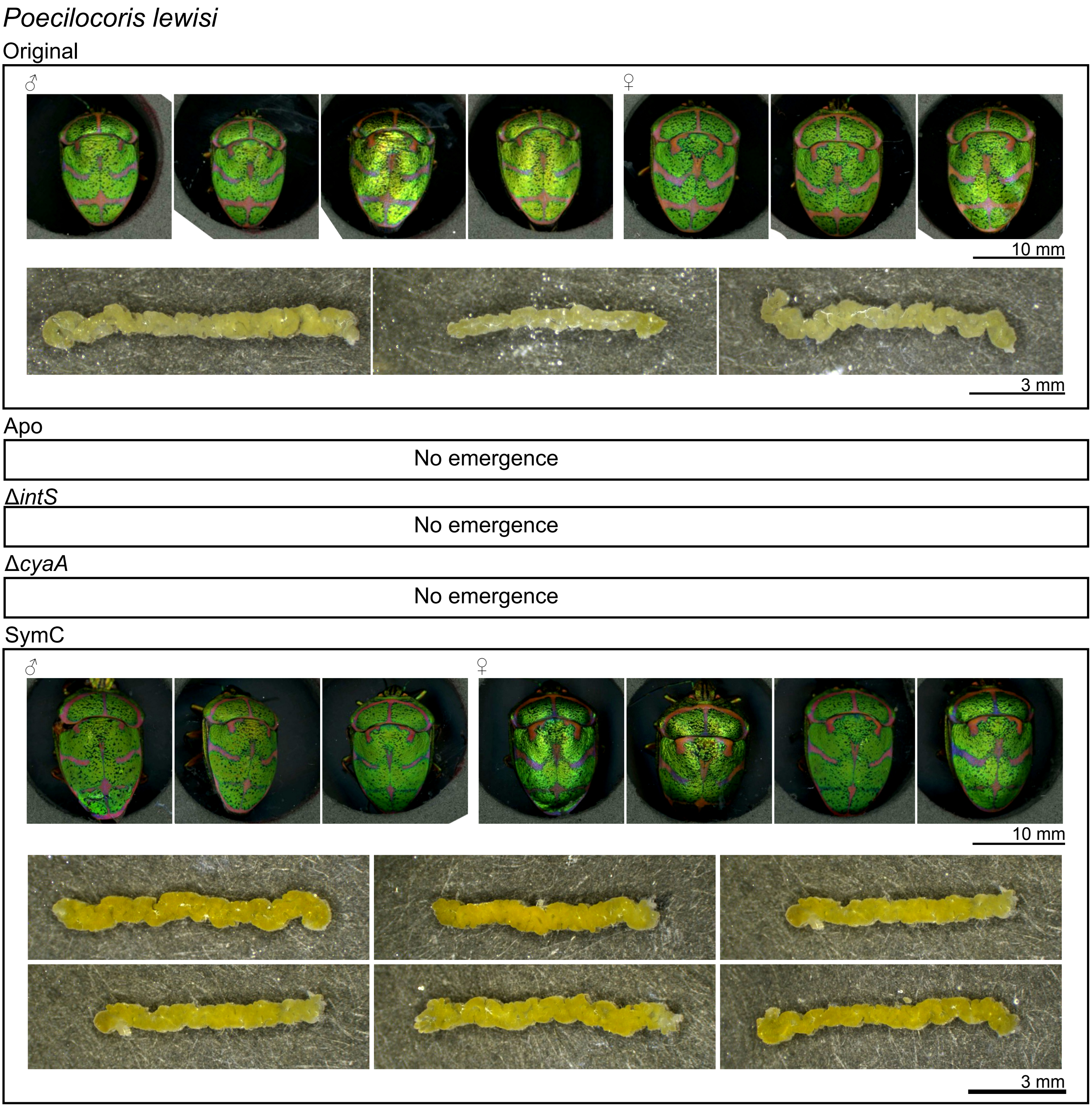


**FIG S7** Adult insects of *P. lewisi* and their symbiotic organs obtained in this study. Also see Fig. 7.

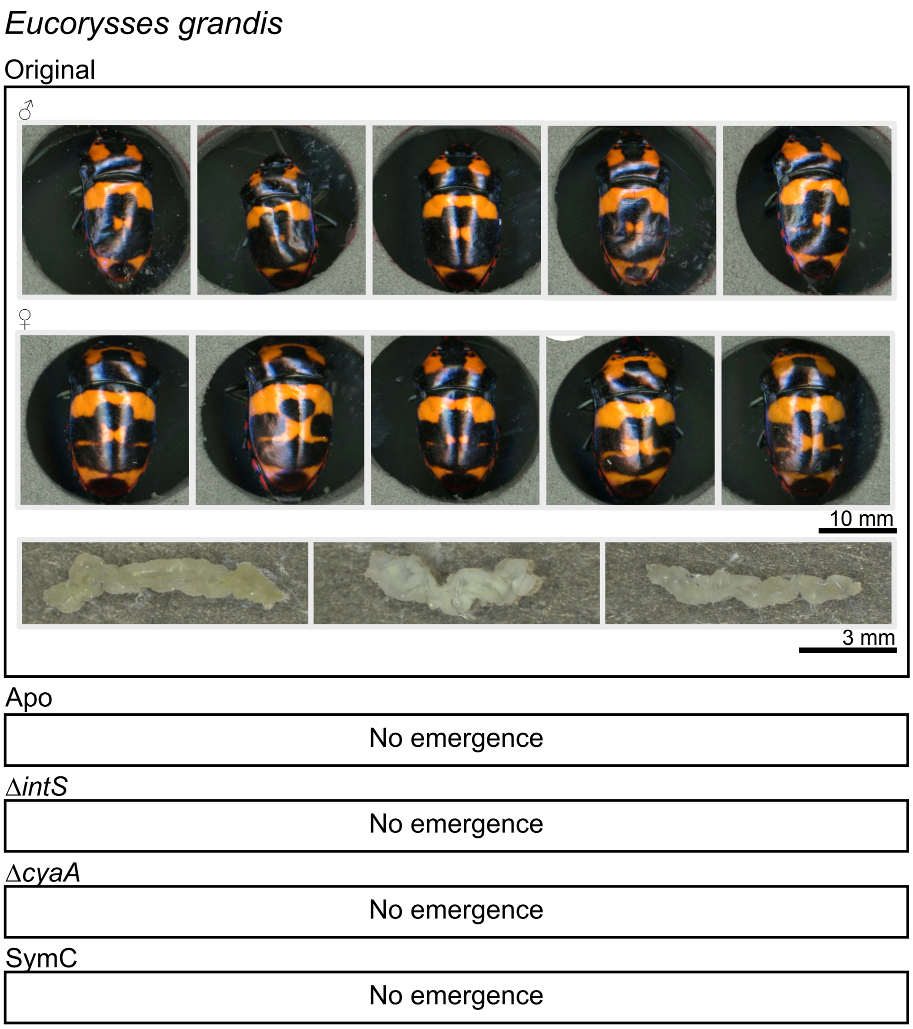


**FIG S8** Adult insects of *E. grandis* and their symbiotic organs obtained in this study. Also see Fig. 8.

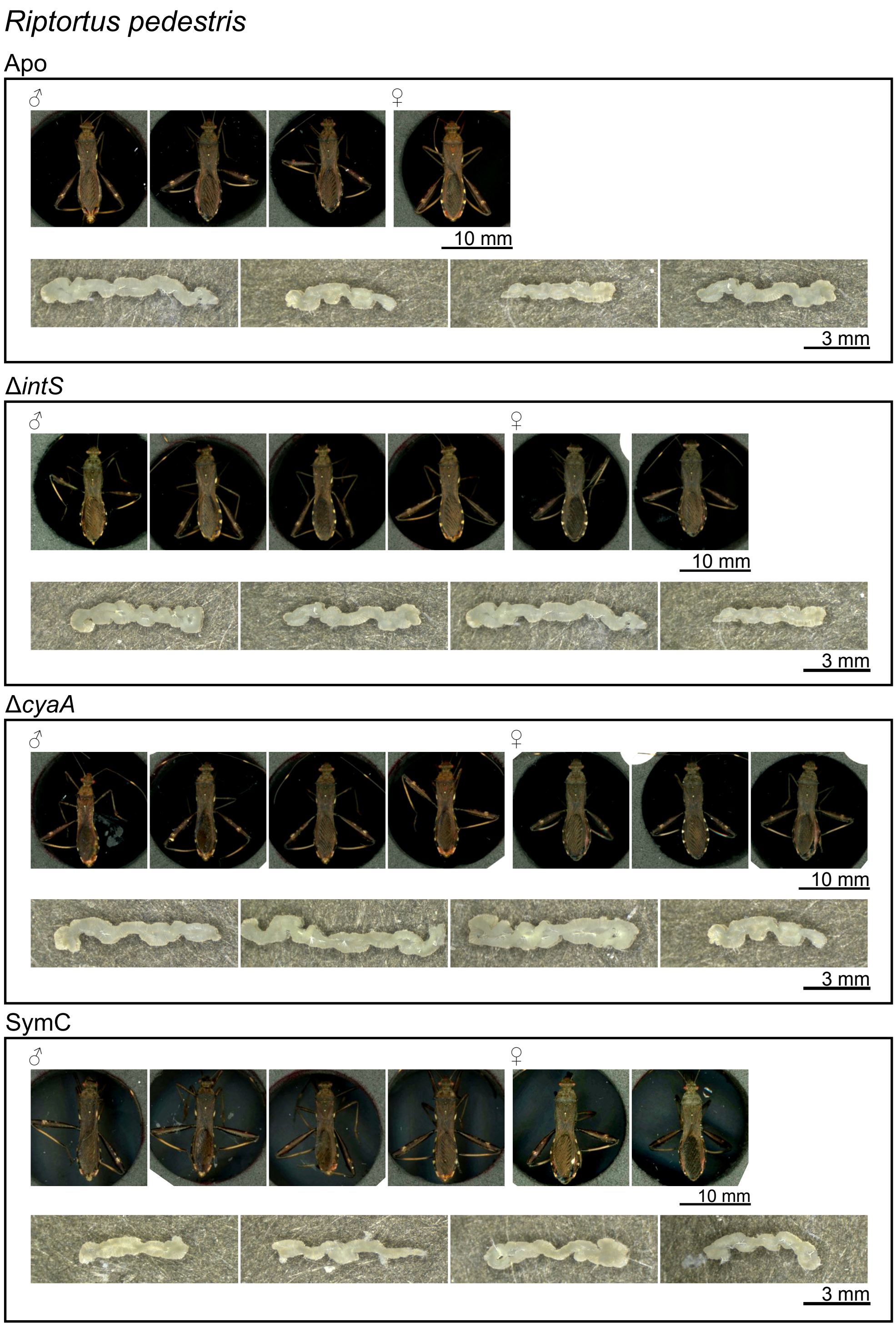


**FIG S9** Adult insects of *R. pedestris* and their symbiotic organs obtained in this study. Also see Fig. 9.

**Table S1**. Laboratory strains of stinkbugs used in this study.

| Insect | Symbiont | Symbiont reference | Collection locality | Collection year | Collector | Rearing condition | Food | Rearing reference | Rearing system | 2nd instar infection check | Adult morphometry and infection check |
|---|---|---|---|---|---|---|---|---|---|---|---|
| **Family Pentatomidae** | | | | | | | | | | | |
| *Plautia stali* | Gammaproteobacteria Enterobacteriaceae *Pantoea* sp. A (Sym A) | Hosokawa et al. (2016a) | Tsukuba, Ibaraki, Japan | 2012 | Minoru Moriyama | 16 h L, 8 h D, 25°C | Raw peanuts; Water with 0.05% vitamin C | Hosokawa et al. (2016a) | 12-15 eggs per plastic cup | 3 days after molting | 41-64 days after hatch |
| *Glaucias subpunctatus* | Gammaproteobacteria Enterobacteriaceae Undescribed | Hosokawa et al. (2016b) | Tsukuba, Ibaraki, Japan | 2018 | Minoru Moriyama | 16 h L, 8 h D, 25°C | Raw peanuts & raw almonds; Water with 0.05% vitamin C | This study | 14-38 eggs per plastic cup | 3 days after molting | 40-67 days after hatch |
| *Nezara viridula* | Gammaproteobacteria Enterobacteriaceae Undescribed | Tada et al. (2011) | Kyoto, Kyoto, Japan | 2019 | Kaoru Ishida & Hideharu Numata | 16 h L, 8 h D, 25°C | Raw peanuts & soybean seeds; Water with 0.05% vitamin C | Tada et al. (2011) | 38-88 eggs per plastic cup | 3 days after molting | 45-67 days after hatch |
| *Halyomorpha halys* | Gammaproteobacteria Enterobacteriaceae Candidatus *Pantoea cerbekii* | Hosokawa et al. (2016b) | Tsukuba, Ibaraki, Japan | 2017 | Minoru Moriyama | 16 h L, 8 h D, 25°C | Raw peanuts; Water with 0.05% vitamin C | This study | 16-29 eggs per plastic cup | 3 days after molting | 56-101 days after hatch |
| **Family Scutelleridae** | | | | | | | | | | | |
| *Lampromicra miyakonus* | Gammaproteobacteria Enterobacteriaceae *Pantoea* sp. C (Sym C) | Hosokawa et al. (2016a) | Ishigaki, Okinawa, Japan | 2020 | Minoru Moriyama | 16 h L, 8 h D, 25°C | Raw almonds & raw cashew nuts; Water with 0.05% vitamin C | Hosokawa et al. (2016a) | 8-15 eggs per plastic cup | 3 days after molting | 45-48 days after hatch |
| *Poecilocoris lewisi* | Gammaproteobacteria Enterobacteriaceae Undescribed | Hosokawa et al. (2019) | Ishigaki, Okinawa, Japan | 2020 | Minoru Moriyama | 16 h L, 8 h D, 25°C | Raw almonds, raw cashew nuts Water with 0.05% vitamin C | This study | 10-13 eggs per plastic cup | 3 days after molting | 50-76 days after hatch |
| *Eucorysses grandis* | Gammaproteobacteria Enterobacteriaceae Undescribed | Hosokawa et al. (2019) | Kamogawa, Chiba, Japan | 2021 | Bin Hirota | 16 h L, 8 h D, 25°C | Raw almonds, raw cashew nuts Water with 0.05% vitamin C | This study | 4-71 eggs per plastic cup | 3 days after molting | 60-76 days after hatch |
| **Family Alydidae** | | | | | | | | | | | |
| *Riptortus pedestris* | Betaproteobacteria Burkholderiaceae *Caballeronia insecticola* | Kikuchi et al. (2005) | Ishigaki, Okinawa, Japan | 2021 | Harumi Yamazaki | 16 h L, 8 h D, 25°C | Soybean seeds Water with 0.05% vitamin C | Kikuchi et al. (2005) | 3-20 eggs per plastic cup | 3 days after molting | 40-43 days after hatch |

---------------
